# Reducing tau ameliorates behavioural and transcriptional deficits in a novel model of Alzheimer’s disease

**DOI:** 10.1101/393405

**Authors:** Eleanor K Pickett, Abigail G Herrmann, Jamie McQueen, Kimberly Abt, Owen Dando, Jane Tulloch, Pooja Jain, Sophie Dunnett, Sadaf Sohrabi, Maria Fjeldstad, Will Calkin, Leo Murison, Rosemary Jackson, Makis Tzioras, Anna Stevenson, Marie D’Orange, Monique Hooley, Caitlin Davies, Iris Oren, Jamie Rose, Chris-Anne McKenzie, Elizabeth Allison, Colin Smith, Oliver Hardt, Christopher M Henstridge, Giles Hardingham, Tara L. Spires-Jones

## Abstract

**Summary:** One of the key knowledge gaps blocking development of effective therapeutics for Alzheimer’s disease (AD) is the lack of understanding of how amyloid beta (Aβ) and tau cooperate in causing disease phenotypes. Within a mouse tau deficient background, we probed the molecular, cellular and behavioural disruption triggered by wild-type human tau’s influence on human Aβ-induced pathology. We find that Aβ and tau work cooperatively to cause a hyperactivity phenotype and to cause downregulation of gene transcription including many involved in synaptic function. In both our mouse model and in human post-mortem tissue, we observe accumulation of pathological tau in synapses, supporting the potential importance of synaptic tau. Importantly, tau depletion in the mice, initiated after behavioural deficits emerge, was found to correct behavioural deficits, reduce synaptic tau levels, and substantially reverse transcriptional perturbations, suggesting that lowering tau levels, particularly at the synapse, may be beneficial in AD.

**Highlights:**

- Expression of human familial Alzheimer’s associated mutant amyloid precursor protein and presenillin 1 with wild-type human tau in the absence of endogenous tau in a novel MAPT-AD mouse model results in behavioural deficits and downregulation of genes involved in synaptic function.
- Tau is present in pre and postsynaptic terminals in MAPT-AD mice and human AD brain. In mice, lowering synaptic tau levels was associated with improved cognition and recovered gene expression.
- These data suggest that Aβ and tau act cooperatively in impairing synaptic function and that lowering tau at synapses could be a beneficial therapeutic approach in AD.

## Introduction

Over 50 million people are living with dementia today, and approximately $800 billion per year is spent worldwide on their health and social care (Prince et al., 2015). Alzheimer’s disease (AD) is the most common cause of dementia, and current drug treatments are only minimally effective and do not prevent brain degeneration or cognitive decline. AD is defined pathologically by the accumulation of amyloid plaques made of aggregated Aβ, neurofibrillary tangles which are intraneuronal deposits of hyperphosphorylated tau protein, and brain atrophy due to neuron and synapse loss. The predominating hypothesis in the AD field, the amyloid cascade hypothesis, posits that changes in amyloid beta (Aβ) initiate the disease (Hardy and Higgins, 1992). Recently, a more nuanced revision of the hypothesis has emerged that views soluble forms of Aβ as the initiator of the cascade and tau as the effector of degeneration in AD (Hyman, 2011). Further, GWAS studies indicate that changes in the innate immune system are important in conferring disease risk (De Strooper and Karran, 2016). How Aβ leads to downstream tau pathology which is associated synapse and neuron degeneration and the role inflammation and the innate immune system plays in this cascade remain key knowledge gaps in the field. We propose that amyloid beta and tau act together both in causing synapse dysfunction/degeneration and in causing neuroinflammation.

Synapses are an important target to study in AD as synapse degeneration is the strongest correlate of cognitive decline (Terry et al., 1991) and synapses are important in disease pathogenesis and the spread of pathological proteins through the brain (Spires-Jones et al., 2017; Spires-Jones and Hyman, 2014). Substantial amounts of evidence implicate oligomeric Aβ in synapse degeneration in model systems and in human post-mortem tissue (Klein, 2013; Koffie et al., 2012; Koffie et al., 2009; Li et al., 2009; Mucke and Selkoe, 2012; Spires et al., 2005; Spires-Jones et al., 2007; Spires-Jones et al., 2009). Some of the toxic effects of Aβ appear to be mediated by cascades which are normally involved in the innate immune system including complement and TREM2 (Hong et al., 2016; Jay et al., 2017; Yeh et al., 2016). Pathological forms of tau are also sufficient to induce synapse loss and circuit dysfunction in models of tauopathy (Crimins et al., 2013; Fox et al., 2011; Hoover et al., 2010; Kopeikina et al., 2012; Menkes-Caspi et al., 2015; Zhou et al., 2017). Recently, immune/inflammatory gene changes have also been observed to contribute to tau toxicity (Leyns et al., 2017; Shi et al., 2017b).

There is accumulating evidence that Aβ and tau may act synergistically to cause synapse and neural circuit degeneration (Ittner et al., 2010; Jackson et al., 2016; Roberson et al., 2011; Shipton et al., 2011; Vargas-Caballero et al., 2017). However, much of the previous work was confounded by the complex differences between mouse and human tau and the inability to precisely control tau expression. To overcome these limitations, we have designed a new model lacking endogenous mouse tau (MAPTnull) and expressing both the APP/PS1 transgene, which causes well-characterized plaque-associated synapse loss (Jankowsky et al., 2004; Koffie et al., 2009), and the rTg21221 line which reversibly expresses wild-type human tau under the control of an inducible promotor (Hoover et al., 2010). This new **M**APTnull+**A**PP/**P**S1+r**T**g21221 AD model (MAPT-AD) allows control over tau levels by suppression of tau transgene expression with doxycycline.

Here we examined the behaviour, pathology, synaptic plasticity, synapse degeneration, transcriptional changes and accumulation of Aβ and tau at synapses in this new model and compared these data to observations of synapses in human post-mortem brain using the high resolution array tomography imaging technique (Kay et al., 2013). MAPT-AD mice develop an age-related hyperactivity phenotype during ageing along with increased expression of genes involved in the innate immune system and decreased expression of genes involved in synaptic function. Pathologically, we observe tau in both pre and post synapses in both human brain and in our MAPT-AD model. Tau was very rarely colocalised with Aβ within individual synapses. Lowering tau levels with doxycycline in the MAPT-AD model reduced synaptic tau levels and ameliorated the behavioural and gene expression phenotypes. Together, these data support the hypothesis that Aβ and tau act together to cause synapse dysfunction. However, this interaction is not likely to be due to physical co-localization of these pathological proteins within the same synapses, at least within the limits of our detection. The presence of tau in synapses appears to mediate toxicity and lowering synaptic tau levels is thus a promising therapeutic target.

## Results

### MAPT-AD mice develop an age-related behavioural phenotype that recovers with tau reduction

The MAPT-AD line is a novel model of AD that combines human mutant APP and PS1 expression (APP/PS1) with regulatable human wild-type tau expression (rTg21221) without the presence of endogenous mouse tau (MAPTnull) (Figure 1A). To understand the effects of combining plaque pathology with human tau expression, we examined pathology and behaviour during ageing in MAPT-AD mice and 3 littermate control genotypes: control (MAPTnull), APP/PS1 only (MAPTnullxAPP/PS1), and human tau only (MAPTnullxrTg21221, Figrure 1B). MAPT-AD mice develop progressive amyloid plaque pathology in the absence of tau pathology (Figure 1C-E). In both genotypes of mice expressing the APP/PS1 transgene, amyloid plaques begin to appear in cortex and hippocampus by 6 months of age and plaque burden increases with age. Plaque deposition differs between APP/PS1 mice and MAPT-AD mice with surprisingly lower plaque burden and smaller individual Thioflavin S positive plaques in mice expressing human tau (Figure 1D, Supplemental Figure 1). Although human tau mRNA and protein could be detected in the two genotypes expressing both the human tau responder gene and the CkTta activator transgene, no tau pathology was observed at any age with staining for phosphorylated or misfolded tau epitopes (AT8, PHF1, Alz50) or with histological staining of fibrils with thioflavin S (Figure 1E). The efficacy of tau staining was confirmed using rTg4510 mouse brain sections (which express a form of tau associated with frontotemporal dementia and develop tangle pathology), verifying that all tau antibodies stained neurofibrillary tangles. MAPT-AD mice did not exhibit age-related atrophy in cortex or hippocampus (Figure S1).

**Figure 1.**
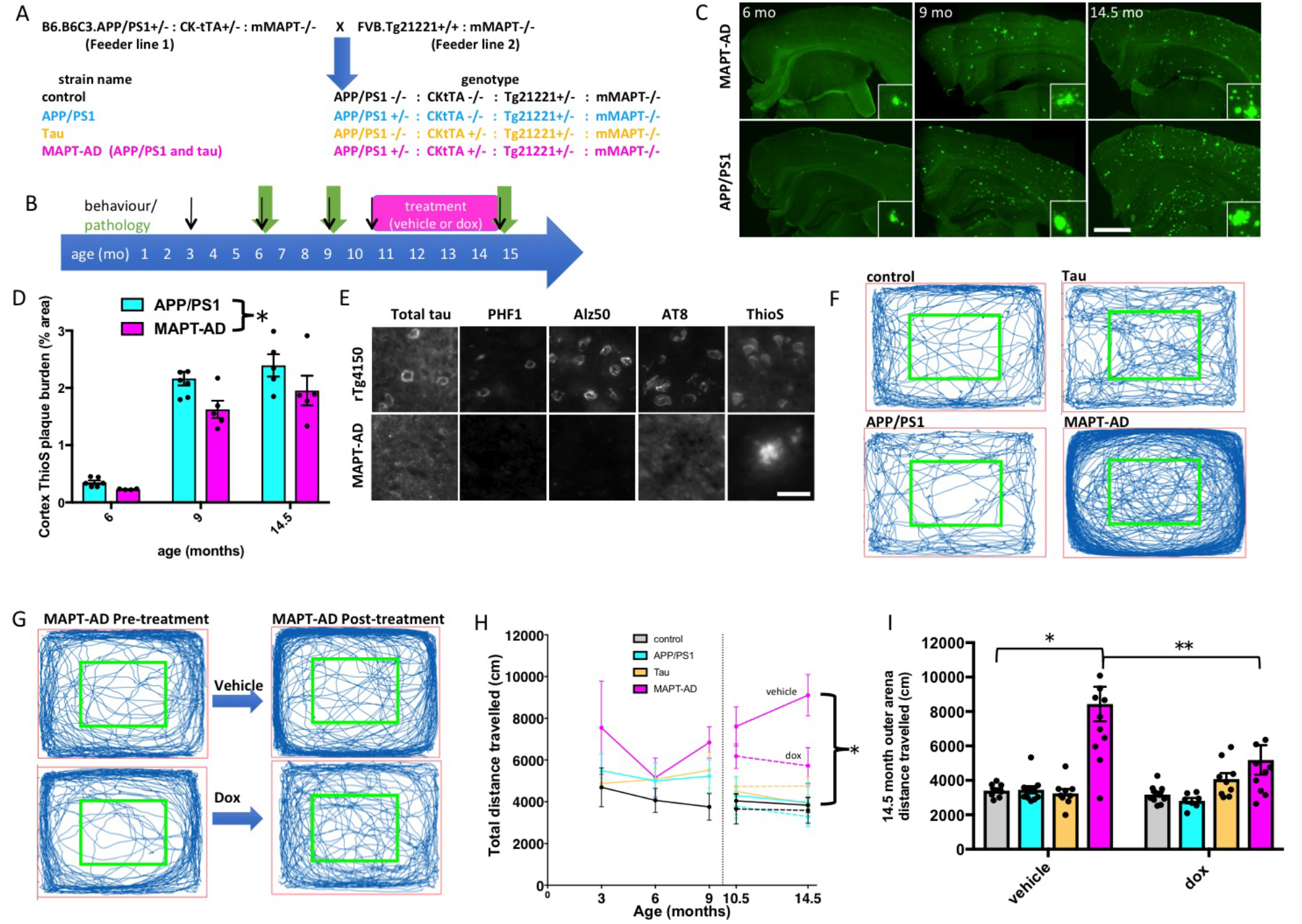
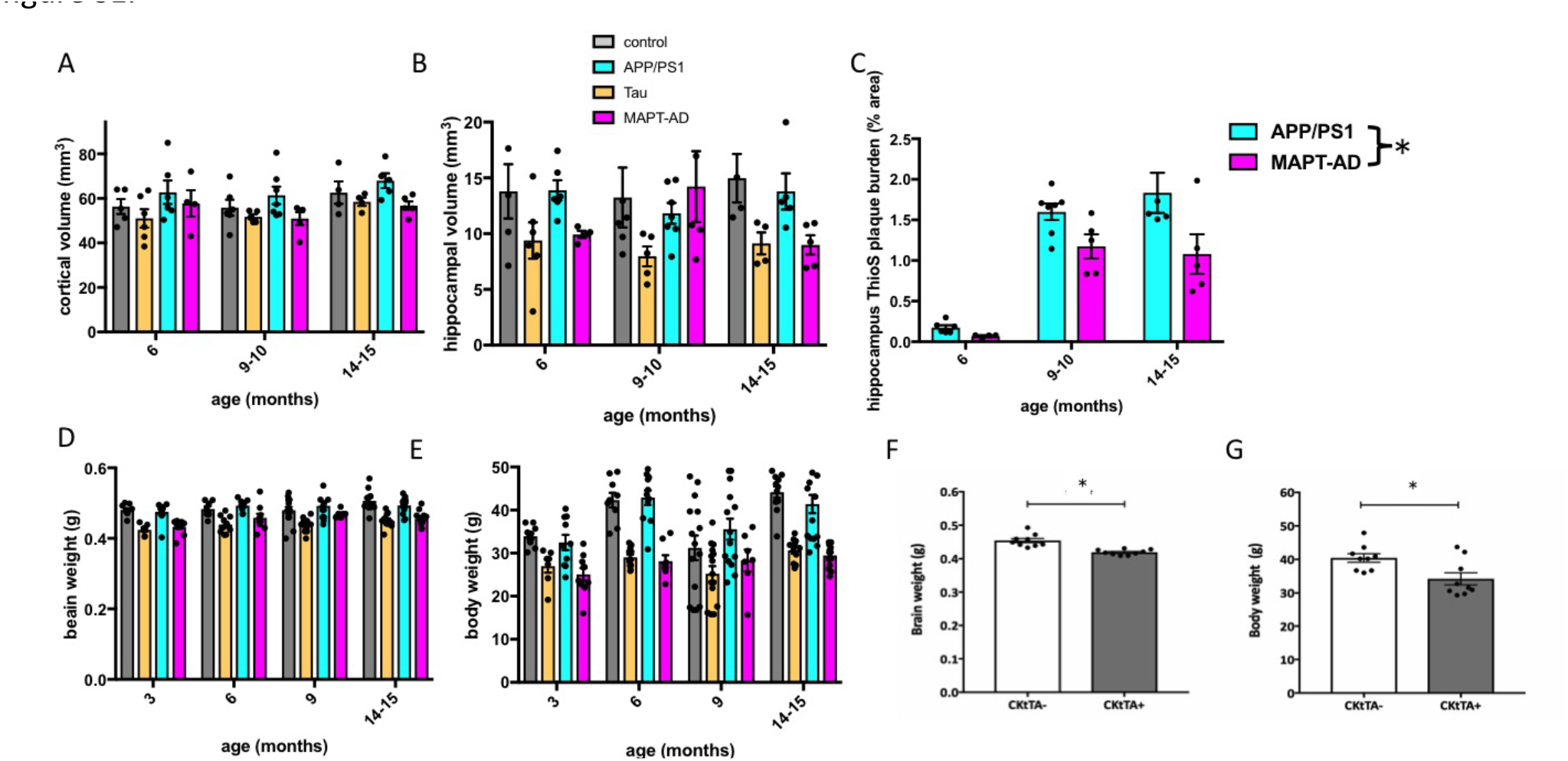
Progressive plaque pathology and reversible hyperactivity phenotype in MAPT-AD mice. The MAPT-AD mouse model was generated by breeding two feeder lines to produce four experimental genotypes of F1 littermates on a consistent outbred strain background (A). Behavior, pathology, and recovery with tau transgene suppression were characterized over time (B). Staining with ThioflavinS (C) shows progressive plaque accumulation in MAPT-AD and APP/PS1 mice. MAPT-AD mice have significantly lower cortical plaque burden that APP/PS1 mice (D, 2 way ANOVA effect of genotype F[2,26]=8.454, p=0.007). Tau is present in MAPT-AD mice but tau pathology does not accumulate in cell bodies or in dystrophic neurites around plaques as shown by staining with total tau, phospho-tau (PHF1 and AT8), or misfolded tau (Alz50) antibodies, which all label tangle pathology in rTg4510 positive control sections (E). The open field test was used as a measure of spontaneous activity. Representative traces from a mouse from each genotype at 10.5 months of age (F), demonstrate the excess activity of the MAPT-AD mice compared to the other three genotypes. This hyperactivity phenotype recovers with dox treatment as seen in representative traces from a MAPT-AD mouse treated with vehicle and one treated with doxycycline and the trace from the same mice after treatment (G) and in the quantification of open field activity (H, repeated measures ANOVA effect of genotype F[3,69]=24.117, p<0.0001; treatment F[1,69]=4.17, p=0.045, interaction F[3,69]=3.58, p=0.018; * Tukey’s multiple comparisons tests vehicle treated MAPT-AD mice are significantly different from control vehicle treated mice (p<0.0001). At 14.5 months of age, MAPT-AD mice travel over 2 times farther in the outer portion of the arena (outside the green box, G) than other genotypes, a phenotype which recovers with dox treatment (2-way ANOVA genotype F[3,69]=19.548, p<0.0001; treatment F[1,69]=3.9990, p=0.0497, interaction F[2,69]=4.770, p=0.004. *, ** Tukey’s multiple comparisons tests p=0.002, p<0.0001). Data shown are means +/- standard error. Dots on bar graphs represent means of individual animals. Scale bars represent 1mm (C, insets 100x100 um), 30 um (E). The dotted lines in H and J represent doxycycline treatment to suppress tau expression from 10.5 – 14.5 months, and the break in the lines indicates a different cohort of mice was used for the 3-6-9 months study and the 10.5-14.5 month study. See also figure S1.

In addition to plaque accumulation and human tau expression, MAPT-AD mice exhibit an age-related hyperactivity phenotype (figure 1F-I). In a cohort of mice tested at 3, 6, and 9 months of age, there was a significant effect of both genotype and age on the distance travelled in the open field (repeated measures ANOVA effect of age F[2,60]=3.5, p=0.038, effect of genotype F[3,30]=3.10, p=0.041). Post-hoc testing revealed a significant increase in open field distance travelled by MAPT-AD mice compared to control mice (Tukey’s multiple comparisons test p=0.026). To test the role of tau in this hyperactivity phenotype, another cohort of mice was aged to 10.5 months and half were treated with doxycycline (dox) from 10.5 to 14.5 months to suppress tau transgene expression. Before treatment, MAPT-AD mice in this cohort travelled significantly further in the open field than the other genotypes (Figure 1F-H, ANOVA effect of genotype F[3,79]=16.87, p<0.0001; Post-hoc Tukey’s multiple comparisons tests show significant increase in the total distance travelled by MAPT-AD mice compared to each of the other genotypes). Treatment with doxycycline for 4 months ameliorates the hyperactivity phenotype in MAPT-AD mice bringing the distance travelled by these mice back to similar levels as all control genotypes (Figure 1G, H). These data suggest vehicle treated mice with both amyloid-beta and tau exhibit increased activity in the open field, which recovers with reduction of tau levels.

To examine whether the increased distance travelled by MAPT-AD mice was due to anxiety, we examined the distance travelled in the inner versus outer portions of the arena. Mice of all genotypes spend approximately 10 times more time in the outer than inner arena indicating a typical avoidance of open areas (supplemental Figure 1). At 14.5 months of age (after treatment), there was no significant effect of genotype or treatment on distance travelled in the inner arena (2-way ANOVA genotype F[3,69]=1.854, treatment F[1,76]=0.204, interaction F[2,69]=0.153, p>0.05). In the outer arena, there were significant effects of genotype, treatment, and an interaction between genotype and treatment on distance travelled (Fig 1I). MAPT-AD vehicle treated mice travelled significantly further in the outer arena than all other groups (Fig 1I). This indicates a potential anxiety phenotype as well as hyperactivity, which recovers with doxycycline treatment

### Tau in synapses may mediate cognitive impairment

To examine the brain changes underpinning the recovery of behaviour with tau suppression, postmortem studies of pathological and molecular changes were carried out in the cohort of mice that had undergone treatment. Treatment of MAPT-AD mice with doxycycline lowered tau mRNA levels by 65% in both MAPT-AD mice and Tau littermates (Figure 2A). APP mRNA levels were 30% higher in MAPT-AD mice than APP/PS1 mice and recovered to normal levels with dox treatment (Figure 2B). In contrast to recovery of APP mRNA, amyloid plaque pathology is unchanged with tau suppression (Figure 2C). The ThioS plaque burden, cross sectional area of individual ThioS stained plaques, AW7 immunostained plaques (which label both the dense core and oligomeric halo surrounding the core), and the area of the oligomeric Aβ halo surrounding plaques were all unchanged with dox treatment. These data indicate that the behavioural recovery was not mediated by reducing amyloid plaque pathology.

**Figure 2:**
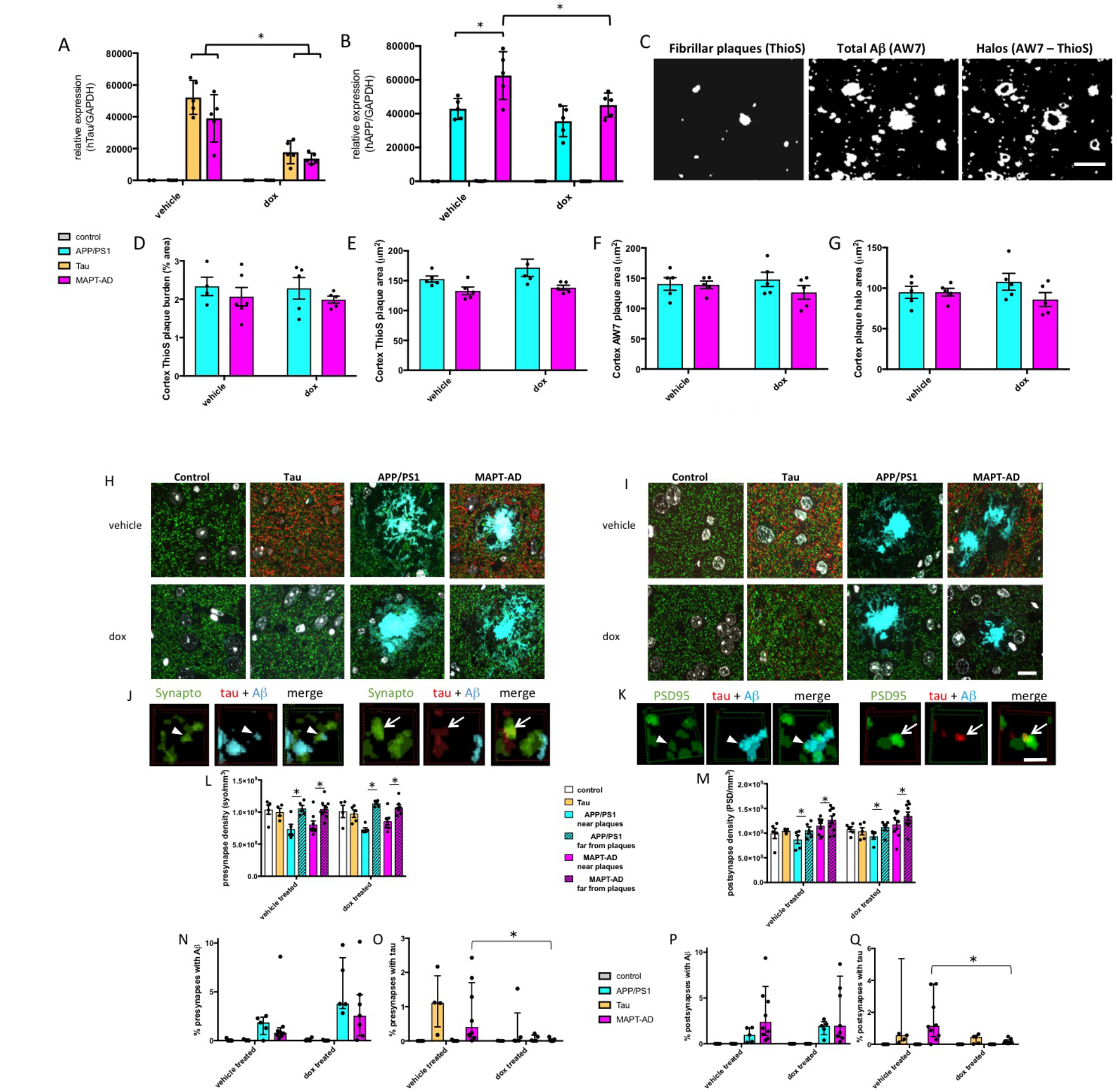
Tau suppression reduces synaptic accumulation of tau. Tau transgene suppression with dox treament reduced tau mRNA levels by approximately 65% as measured by qPCR (A, * 2-way ANOVA effect of treatment F[1,31]=42.22, p<0.0001). APP mRNA levels (B) were increased by 30% in MAPT-AD mice, an effect which was ameliorated by dox treatment (2-way ANOVA genotype F[3,31]=153.5, p<0.0001, treatment F[1,31]=7.912, p=0.0084, interaction F[3,31]=3.468, p=0.0279, * post-hoc Tukey’s test p<0.01). Tau suppression did not change amyloid pathology in MAPT-AD mice. Fibrillar plaques were measured with ThioS, total Ab with AW7 immunostaining, and oligomeric Ab halos were measured by subtracting the fibrillar cores from total Ab staining (G). To investigate synapse loss and synaptic proteins, array tomography ribbons were stained for presynaptic terminals (H, synaptophysin, green), or postsynaptic densities (I, PSD, green) along with human tau (red), and amyloid beta (AW7, cyan). Maximum intensity projections of 10 serial 70 nm sections are shown in H and I. Three-dimensional reconstructions of 5 consecutive serial sections from processed image stacks of a MAPT-AD mouse demonstrate presynaptic (J) and postsynaptic (K) terminals positive for tau (arrows) or Aβ (arrowheads).

Synapse density around plaques and the accumulation of synaptic Aβ and tau were determined using array tomography. More than 673,000 postsynaptic densities labelled with PSD95 and 415,000 presynaptic terminals labelled with synaptophysin were analysed from cortical samples from 4-11 mice per group (average 13,000 PSDs and 9,655 presynaptic puncta per mouse). Density of both synaptophysin (figure 2L) and PSD95 (figure 2M) labelled puncta was decreased near plaques in both genotypes that have plaques (APP/PS1 and MAPT-AD mice, 3-way ANOVA effect of plaque distance synaptophysin F[1,42]=60.49, p<0.0001, PSD95 F[1,50]=8.15, p=0.006). Treatment with doxycycline to reduce tau levels did not prevent this plaque-associated synapse loss. The density of pre and post synapses near plaques was not significantly different between treatment groups or genotypes. Oligomeric Aβ accumulated in a subset of synapses near plaques in both APP/PS1 and MAPT-AD mice (median presynaptic terminals near plaques containing Aβ 1.9% in APP/PS1 mice, 0.8% in MAPT-AD mice, median postsynaptic terminals 1.0% in APP/PS1 mice, 2.4% in MAPT-AD mice). The percentage of both pre and postsynaptic terminals containing Aβ was higher near plaques than far from in both APP/PS1 and MAPT-AD mice (independent samples Mann-Whitney U test PSD95 p<0.01, synaptophysin p<0.0001 for all groups near vs far from plaques). The percentage of synapses containing Aβ near plaques was not different between APP/PS1 and MAPT-AD mice (independent samples Mann-Whitney U test p>0.05). There was also no effect of lowering tau levels on accumulation of Aβ in synapses near plaques (independent samples effect of genotype, Mann-Whitney U test p>0.05).

Tau was detected in median of 1.2% of PSDs (figure 2O) and 0.4% of presynapses (figure 2Q) in vehicle treated MAPT-AD mice and 0.6% of PSDs and 1.1% of presynapses in vehicle treated Tau mice (the only genotypes expressing tau). Unlike Aβ, the percentage of synapses containing tau was not different near plaques in the MAPT-AD group. The percentage of synapses containing tau were significantly different between genotypes (independent samples Mann-Whitney U test for genotype p<0.0001). Doxycycline treatment significantly lowered synaptic tau levels only in the MAPT-AD group (data split by genotype, effect of treatment independent samples Mann-Whitney U test p=0.004 for PSD95, p=0.004 synaptophysin). The approximate 30-fold reduction in presynaptic and 8-fold reduction in postsynaptic tau levels in MAPT-AD mice may contribute to the improved hyperactivity phenotype observed in mice treated with doxycycline. Only very rare PSDs stained for both Aβ and tau (<0.006% of pre <0.005% post synapses in vehicle treated MAPT-AD mice).

Quantification reveals significant pre (L) and post (M) synapse loss near plaques in APP/PS1 and MAPT-AD mice which is not rescued by lowering tau levels with doxycycline (dox) treatment. The percentage of presynapses (N) and postsynapses (P) positive for Aβ is not different between MAPTnullxAPP/PS1 mice and MAPT-AD mice, nor is it affected by dox treatment. The percentage of presynapses (O) and postsynapses (Q) containing tau is significantly lowered by dox treatment in MAPT-AD mice (* Mann-Whitney U test p=0.004). Data represent mean + SEM (L, M) and median + interquartile range (N-Q). Scale bars represent 1 mm in B, 30 μm in C, 10 μm in H and I, 1 μm in J and K.

### Tau suppression reverses transcriptional changes

In addition to targeted studies of synapses, Aβ, and tau postmortem, we performed unbiased RNAseq experiments to look for transcriptional changes in MAPT-AD mice and whether these recover with tau transgene suppression (Raw data available in Supplemental table 2 and ***ADD DOI after acceptance***). MAPT-AD mice had 1531 transcripts that were detected with FPKM >1 which were significantly altered compared to MAPTnull control mice and 127 of these were changed by greater than 2 fold (Figure 3C). The gene changes in MAPT-AD mice compared to control are much larger than either APP/PS1 mice (Figure 3A, 81 transcripts with significant >2 fold change) or Tau mice (Figure 3B, 6 transcripts with significant >2 fold change) compared to controls. Gene Ontology enrichment analysis indicates that the upregulated genes in MAPT-AD mice are dominated by an inflammatory response (including *Trem2, Gfap, Cd68, C1q*, and *H2-Eb1*), while downregulated genes are involved in synaptic function including glutamate signalling (AMPA and NMDA receptor subunits, *Gria2, Gria3, Gria4, Grin2a, Homer2*, and *Camk2b*). One synaptic transcript that was significantly upregulated is cellular prion protein (*Prnp*,), which is very interesting since it is a known synaptic binding partner of Aβ (Um et al., 2012).

**Table 2:**
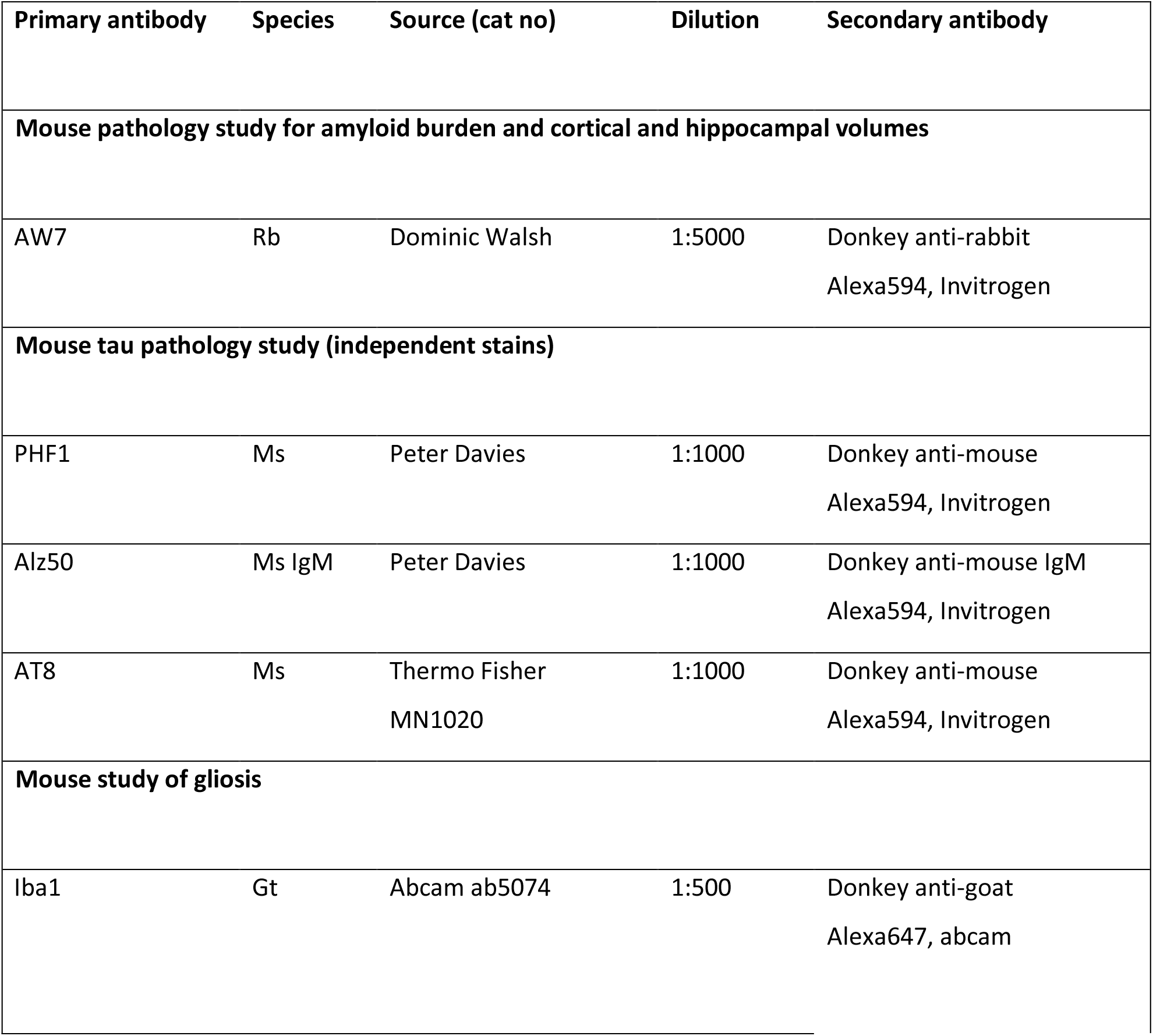
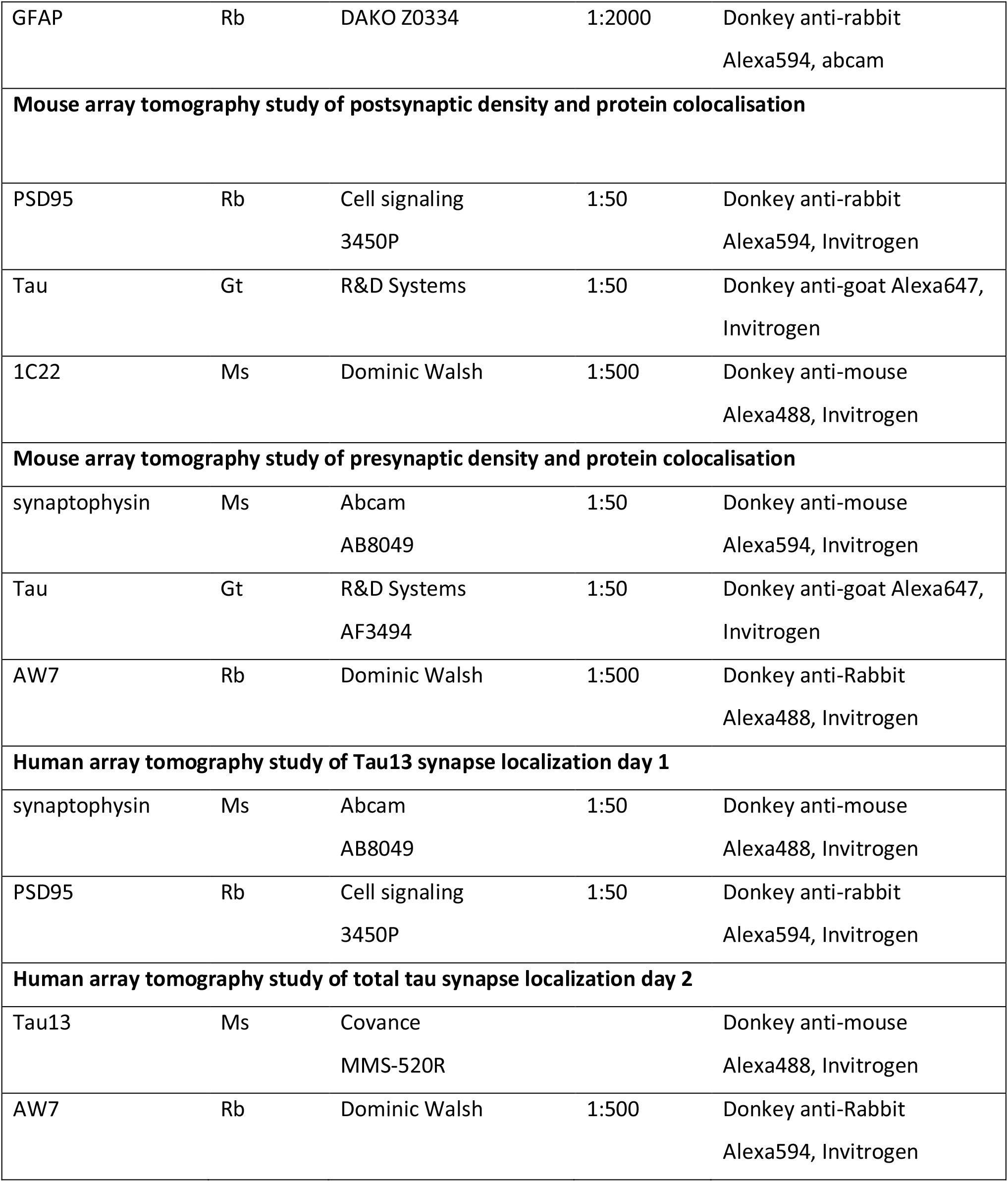
Antibodies used in array tomography and histology studies. Host species were mouse (Ms), Rabbit (Rb), goat (Gt), and guinea pig (Gp).

**Figure 3:**
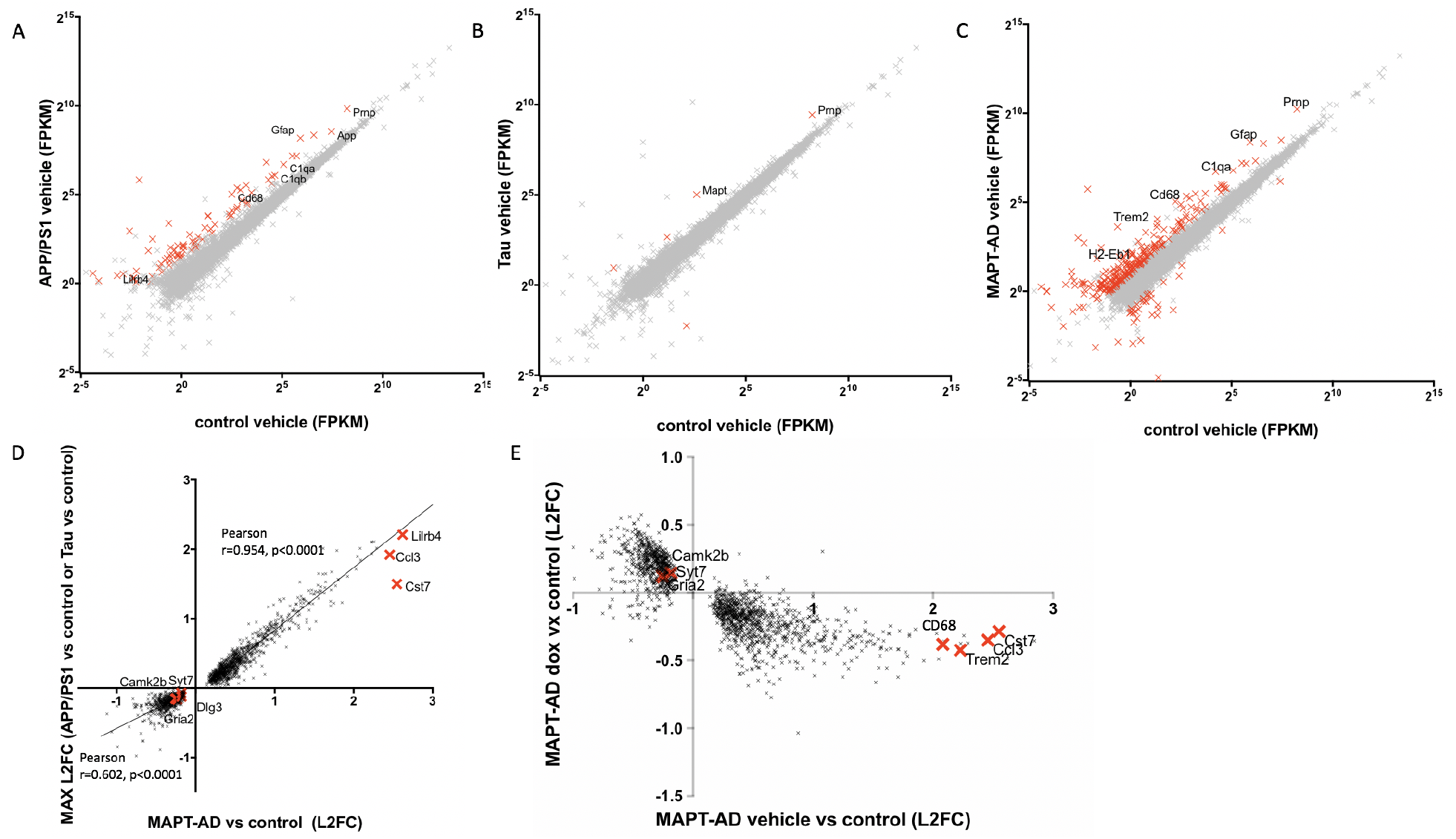
Transcriptional changes in MAPT-AD mice that are reversed by tau suppression. RNAseq of APP/PS1 brain compared to controls reveals significant changes in gene expression (A, FPKM, Fragments Per Kilobase of transcript per Million mapped reads). Wild-type human tau induced changes to a lesser extent (B). MAPT-AD mice (C) had more significant changes than when either APP/PS1 or Tau were expressed on their own. Transcripts changed more than 2 fold with an adjusted p value of p<0.05 are shown in red in panels A-C. A subset of crosses are labelled to show transcripts of interest that are changed in including genes involved in inflammation (Cd68, C1q, Gfap, risk factors Trem2 and histocompatibility class II (H2-eb1), and cellular prion protein (Prnp) which is a synaptic binding partner of Aβ. Examining only the genes significantly chanted in MAPT-AD mice compared to control mice and comparing the log(2) fold change (L2FC) of MAPT-AD mice compared MAPTnull control mice to the maximum L2FC of either MAPTnullxAPP/PS1 or MAPTnullxrTg21221 compared to MAPTnull controls (D) shows that upregulated genes are for the most part not differentially regulated in MAPT-AD mice compared to those expressing APP/PS1 or tau alone (slope 0.90, 95% CI 0.89 to 0.92). Downregulated genes in MAPT-AD mice (D) are differentially regulated in MAPT-AD mice compared to those expressing APP/PS1 or tau alone (slope 0.57, 95% CI 0.51 to 0.63). Red crosses in C show transcripts of interest which are changed more in MAPT-AD mice than in APP/PS1 or Tau mice including downregulated genes involved in synaptic function (Gria2,Camk2b, Dlg3, and Syt7) and upregulated genes involved in inflammation (Lilrb4, Ccl3, and Cst7). Dox treatment to reduce tau levels reverses transcriptional changes in MAPT-AD mice (E, linear regression slope = -0.34, 95% CI -0.36 to -0.33). Each point represents a single transcript. See also related figure S2.

Upregulation of genes in MAPT-AD mice appears to be driven by Aβ and tau independently without an additive effect since the fold induction of upregulated genes is very similar in MAPT-AD mice to the maximum fold induction in either APP/PS1 or Tau mice, illustrated by the fact that most of these genes correlate strongly between MAPT-AD and the maximum fold change in either APP/PS1 or Tau (Figure 3D). There are a few interesting genes upregulated more in MAPT-AD mice than in APP/PS1 or Tau mice including several involved in inflammation (Lilrb4, Ccl3, and Cst7). Cst7 is interesting as it is upregulated in disease associated microglia in APP/PS1 mice (Keren-Shaul et al., 2017). In contrast to the relatively few changes seen in upregulated genes, Aβ and tau act additively in downregulating gene expression. The fold downregulation compared to controls in MAPT-AD mice is more than the maximum change in either APP/PS1 mice or Tau mice (Figure 3D). Doxycycline treatment ameliorates gene expression changes in MAPT-AD mice (Figue 3E) and reverses the mild changes in Tau mice (figure S2), indicating that lowering tau levels protects against gene expression changes.

To test whether the recovery of gene expression with tau suppression was due to a prevention of further changes with age or a recovery of existing changes at the time treatment began, we analysed a subset of transcripts by RT-PCR at 9-10 months of age (an age before treatment started) and validated the RNAseq data again in 14.5 month old brain samples that had been treated with vehicle or dox. The subset of genes tested indicate that the amelioration of gene expression changes with dox was due to a prevention of further worsening and not a recovery (supplemental figure 2). Since many of the upregulated inflammatory genes are expressed in glia, we examined astrocyte and microglial burdens. In agreement with the overall trend observed with RNAseq that upregulated genes are driven independently by Aβ or tau, an increase in gliosis was observed in both genotypes with human Aβ, MAPT-AD and APP/PS1 mice. These did not recover with dox treatment (figure S2).

### Tau is present in pre and post synapses of human AD cases

To confirm the translational relevance of the contribution of synaptic tau to cognitive decline in our new model, we examined the localization of tau and Aβ at synapses in samples of superior temporal gyrus from human AD and control subjects. In total 99,967 postsynapses and 100,012 presynapses form 6 AD and 6 control subjects were examined (mean 8,331 post and 8,334 pre synapses examined per case, data found in supplemental table 3). Cases were stained with the pan-Aβ antibody AW7, a tau antibody, a presynaptic marker, and a postsynaptic marker in a two-day protocol to allow localization of Aβ and tau together within individual pre and postsynapses. As previously reported, Aβ is present in a subset of synapses in AD brain with significantly more positive synapses within 20 μm of a plaque (10.33% PSD and 14.29% synaptophysin puncta positive for Aβ near plaques, <1% PSD or synaptophysin positive for Aβ far from plaques, p<0.05 Independent samples Mann-Whitney U test for both pre and post synapses). In array tomography, the tau13 antibody recognized neurofibrillary pathology but not normal axonal tau, and labelled a small subset of pre and post-synapses (Figure 4). As observed in the mice, tau synaptic localization was not significantly different near versus far from plaques. Also in agreement with the mouse data, only very rare synapses were positive for both tau and Aβ staining (<0.02% on average).

**Figure 4:**
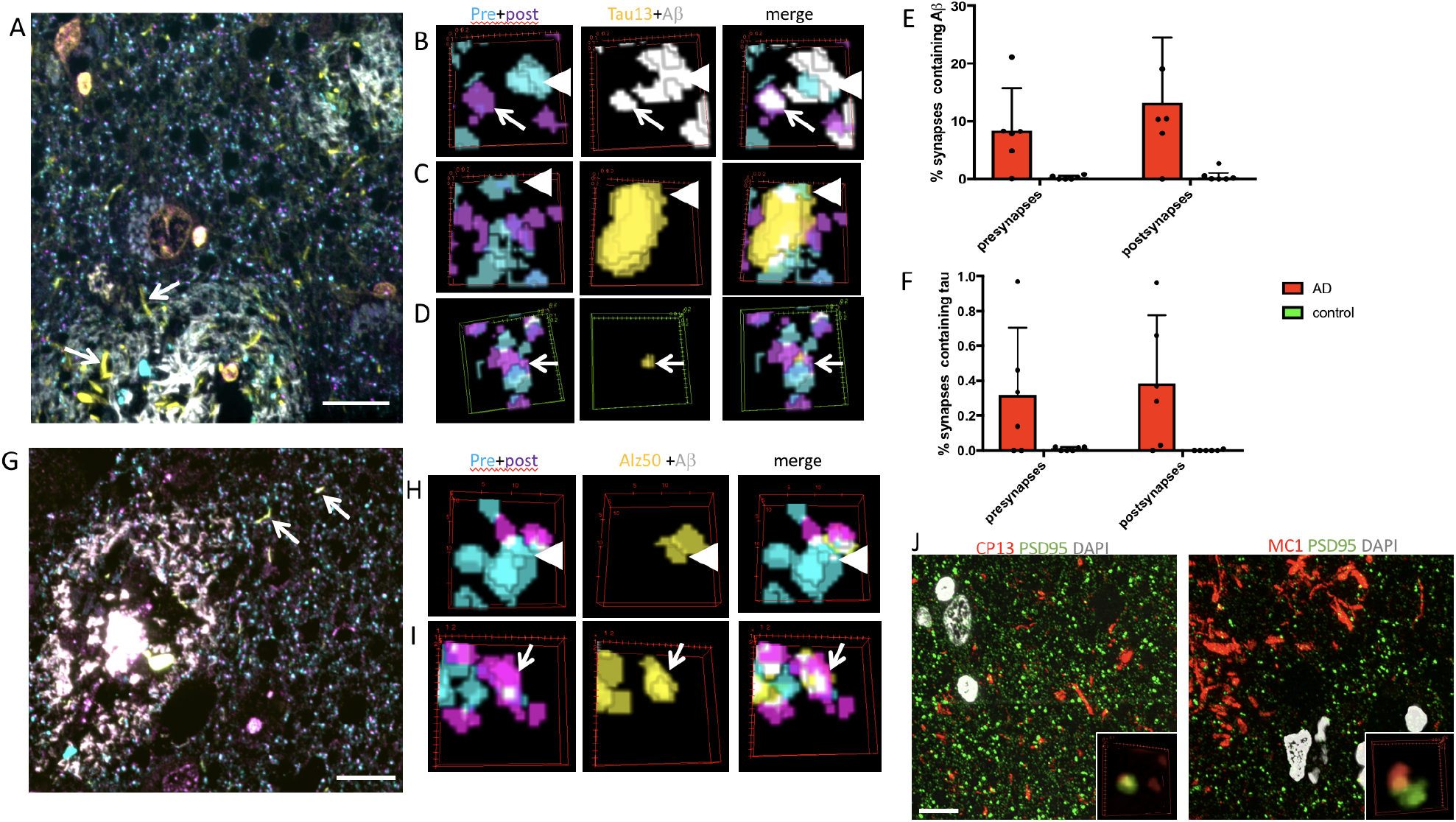
Tau is found in pre and post synapses in human AD brain. Array tomography was used in human postmortem brain tissue to stain Aβ (white), Tau13 (yellow), PSD95 (magenta), and synaptophsyin (cyan) (A-D). Tau13 stains neuropil threads (arrows, A). Examining individual synapses revealed that A β was present in 8.49% of presynaptic terminals (B arrowheads) and 6.98% of postsynaptic densities (B arrows) near plaques in AD cases (B, E). Tau13 staining was observed in 0.32% of presynaptic terminals (arrowheads C, quantified in F), and 0.38% of postsynaptic terminals (arrows, D quantified in F). Misfolded tau labeled with Alz50 (G-I, yellow) was also observed in neuropil threads (arrows G) and in presynapses (arrowheads, H) and post synapses (arrows, I). Tau phosphorylated at serine 202 (labeled with CP13) and misfolded (residues 5-15 near312-322, labelled with MC1) was also observed in PSDs (J). Images in A, G, and large panels are maximum intensity projections of 10 serial sections. Scale bar represents 10 um in A, G, J. B-D, H, I, and insets show three-dimensional reconstructions of a 2 micron by 2 micron region of interest in 5 consecutive serial 70nm sections.

## Discussion

The lack of disease modifying treatments for AD remains a huge unmet clinical need. Advances in understanding of the mechanisms of neurodegeneration will pave the way for developing effective treatments. Synapse degeneration is the strongest pathological correlate of cognitive decline in AD and a potentially important driver of disease pathogenesis. Previous work by our group and others strongly implicated soluble Aβ and tau separately in synapse dysfunction and loss in AD (Klein, 2013; Koffie et al., 2012; Koffie et al., 2009; Kopeikina et al., 2012; Mucke and Selkoe, 2012; Spires-Jones et al., 2017; Spires-Jones and Hyman, 2014). Here we tested the hypothesis that Aβ and tau act together to cause neural circuit dysfunction. Evidence has been growing over the past decade to support this idea both from work showing that lowering tau levels protects against Aβ mediated synaptic plasticity deficits and from studies indicating that dendritic tau mediates Aβ synaptotoxicity (Ittner et al., 2010; Roberson et al., 2011; Roberson et al., 2007; Shipton et al., 2011; Zempel et al., 2010). In the novel MAPT-AD model, we observe an age-related hyperactivity phenotype and downregulation of genes involved in synaptic function. Reducing tau expression levels ameliorated the behavioural phenotype and lowered synaptic tau levels without recovering synapse density around plaques. The potential importance of synaptic tau in human disease was confirmed by quantifying the presence of tau at pre and postsynaptic terminals in AD brain.

Potential molecular mechanisms linking Aβ and tau to synapse and circuit dysfunction include calcium dysregulation and calcineurin activation, which are known to contribute to Aβ toxicity and spine collapse *in vitro*, and *in vivo*, and have recently been linked to tau mediated synapse impairment (Hudry et al., 2012; Kuchibhotla et al., 2008; Mattson et al., 1992; Wu et al., 2010; Yin et al., 2016; Zempel et al., 2010). Abnormal activation of synaptic receptors by Aβ has also been shown to induce activation of kinases including Fyn and GSK3-β which affect tau phosphorylation and synapse collapse (Ittner et al., 2010; Lovestone et al., 2014; Marzo et al., 2016; Purro et al., 2012; Roberson et al., 2011; Sellers et al., 2018; Small and Duff, 2008). Our RNAseq results add to the literature implicating cellular prion protein at the interface between Aβ and tau as increases in PrPc mRNA in MAPT-AD mice was the largest change observed with RNAseq and these levels recover with tau suppression. PrPc has been shown to interact with oligomeric Aβ where it is thought to act via metabotropic glutamate receptor 5 complexes to impair synaptic function (Barry et al., 2011; Haas and Strittmatter, 2016; Hu et al., 2018; Hu et al., 2014; Jarosz-Griffiths et al., 2016). This pathway could involve tau since binding of Aβ to PrPc can activate Fyn and cause tau phosphorylation (Um et al., 2013; Um et al., 2012). While many of the proposed mechanisms of synapse degeneration focus on post-synaptic processes, our data clearly show accumulation of both Aβ and tau in pre as well as postsynaptic terminals. Tau has recently been shown to bind to presynaptic vesicles in human AD and Drosophila models, where it impairs neurotransmitter release (McInnes et al., 2018; Zhou et al., 2017). Similarly, it is also becoming clear that Aβ exerts effects on presynaptic function (Ovsepian et al., 2018).

Our RNAseq results strongly implicate non-neuronal cells as key participants in the interplay between Aβ and tau. TREM2, clusterin, and CD33, genes involved in the innate immune system that have recently been implicated in AD risk by GWAS studies, were elevated in MAPT-AD mice compared to controls. Several members of the complement cascade family were also changed in MAPT-AD mice, which is important due to the recent discovery of complement mediated microglial engulfment of synapses in plaque bearing AD model mice (Hong et al., 2016; Shi et al., 2017a). Our data indicate that beyond contributing to disease risk, presumably through amyloid, the innate immune system is also likely involved in the cascade from amyloid to tau in AD pathogenesis. The gene changes observed in our model indicate that Aβ and tau act cooperatively to cause downregulation of genes and largely independently in gene upregulation. Downregulated genes were predominantly involved in excitatory synaptic function, which is supported by recent data implicating tau in toxicity to excitatory over inhibitory neurons (Fu et al., 2017).

Synapses are highly plastic structures, which have the potential for recovery with interventions. Indeed, most successful drugs used in nervous system disorders act at the synapse; therefore synaptic changes are an obvious target for disease-modifying agents in neurodegenerative disorders. Recent work has focused on removing Aβ from synaptic receptors as a therapeutic avenue. For example, a compound that displaces Aβ from sigma-2 receptors is now in clinical trials (Izzo et al., 2014), clinicaltrials.gov). Our data indicate that lowering pathological tau specifically at synapses may also be an effective therapeutic strategy.

## Acknowledgements

We wish to thank our human brain tissue donors and their families for their valuable contributions to this work. Many thanks also to Prof George Carlson and Ms Rose Pitstick for their input on mouse breeding and providing mouse lines. This work was funded by the UK Dementia Research Institute, the European Research Council (ALZSYN), Alzheimer’s Society, Alzheimer’s Research UK and the Scottish Government Chief Scientist Office (ARUK SPG2013-1), a Wellcome Trust-University of Edinburgh Institutional Strategic Support Fund, MND Scotland, and Alzheimer’s Society. SD contributed to this work funded by an Alzheimer’s Society summer studentship. MF contributed to this work funded by a summer studentship from Medical Research Scotland. TSJ gratefully acknowledges affiliations with the FENS-Kavli Network of Excellence, the Centre for Dementia Prevention, the Euan MacDonald Centre, and Edinburgh Neuroscience.

## Author Contributions

Conceptualization, G.H. and T.L.S-J.; Methodology, E.K.P., J.T., G.A.C., and T.L.S.-J., Software, T.L.S.-J.,and E.A.; Formal Analysis, O.D., M.H., I.O. G.E.H. and T.L.S.-J.; Investigation, E.K.P., J.M., K.A., J.T., A.G.H., O.N., P.J., S.D., S.S.S., A.S., M.F., W.C., L.M., R.J.J., M.T., M.D’O., J.R., R.P., C.-A.M., C.S., and C.M.H; Writing – Original Draft, E.K.P, and T.L.S.-J.; Writing – Reviewing & Editing, C.M.H, G.H., and T.L.S.-J.; Supervision, I.O., O.H., G.H., and T.L.S.-J.; Funding Acquisition G.H., and T.L.S.-J.

## METHODS

### EXPERIMENTAL MODEL AND SUBJECT DETAILS

#### Animals

All animal experiments conformed to national and institutional guidelines including the Animals [Scientific Procedures Act] 1986 (UK), and the Council Directive 2010/63EU of the European Parliament and the Council of 22 September 2010 on the protection of animals used for scientific purposes, and had full Home Office ethical approval. Mice were bred in house and group housed in a 12h/12h light/dark cycle with ad libitum access to food and water. Both sexes of mice were used in all experiments (see supplemental table 1 for details of all mice used including sex, age, and weight information). Littermates were randomly assigned to experimental groups in experiments to reduce tau transgene expression and experimenters were blind to genotype and treatment.

**Table 1:** Human subject characteristics

| case | diagnosis | age | sex |
| --- | --- | --- | --- |
| 1442 | AD | 80 | f |
| 1446 | AD | 84 | m |
| 1547 | AD | 77 | m |
| 1564 | AD | 90 | m |
| AD5 | AD | 75 | f |
| BBN24526 | AD | 79 | m |
| HC | control | 66 | m |
| HC2 | control | 69 | m |
| HC3 | control | 75 | m |
| HC6 | control | 95 | m |
| BBN28406 | control | 79 | m |
| BBN19686 | control | 77 | f |

#### Human subjects

Brain tissue samples were taken from superior temporal gyrus of 6 AD and 6 control subjects in the Edinburgh Sudden Death Brain Bank or the Massachusetts General Hospital Alzheimer’s Disease Research Centre Brain Bank. Characteristics of human subjects can be found in table 1 and synapse data in supplemental table 3. Average age was 81 for AD cases (range 75-90) and 77 for control cases (range 69-95). All AD cases were neuropathologically confirmed and were Braak stage V or VI. Control cases had no neurological phenotype. All human experiments were reviewed and approved by the Sudden Death Brain Bank ethics committee and the ACCORD medical research ethics committee (Academic and Clinical Central Office for Research and Development at the University of Edinburgh and National Health Service Lothian, ethical approval number 15-HV-016).

### METHOD DETAILS

#### Generation of new mouse line

For the new **M**APTnull **A**PP/PS1 r**T**g21221 (MAPT-AD) model line, 4 genotypes were used to compare mice with (1) no transgene expression on a MAPTnull background, (2) mice expressing human familial AD mutant APP and PS1 to generate A β pathology (MAPTnullxAPP/PS1), (3) mice expressing 0N4R wild-type human tau (MAPTnullxrTg21221), and (4) mice expressing both human tau and the APP/PS1 transgene (MAPT-AD, **Figure 1**). All mice were homozygous for deletion of mouse tau and heterozygous for the human wild-type tau transgene which is only expressed when the tetracycline transactivator transgene is also present. All experimental mice were F1 crosses from two feeder lines to maintain a controlled outbred background strain with consistent proportions of B6, B6C3, and FVB backgrounds. Parent strains used to generate the MAPT-AD feeder lines were: (1) B6C3 APP/PS1 mice expressing human APP with the Swedish mutation and human presenilin 1 with an exon 9 deletion under the control of the Thy1 promoter (B6C3-Tg(APPsw,PSEN1dE9)85DboMmjax, Jax 34829, (Jankowsky et al., 2004); (2) MAPTnull mice on the C57BL/6 background strain which have the first exon of the MAPTgene replaced with EGFP (Tucker et al., 2001); (3) mice expressing the tetracycline transactivator under the control of the calcium calmodulin kinase 2 alpha promoter CK-tTA on the C57BL/6 backgrounds strain (B6.Cg-(Camk2a-tTA)1/MmayDboJ ,(Yasuda and Mayford, 2006)); (4) Tg21221 mice expressing human wild type tau under a dox-off tetracycline transactivator promotor (FVB-Tg(tetO-0N4R-MAPTwt)21221, (Hoover et al., 2010)). One feeder line was generated by crossing FVB.MAPTnull mice with the Tg21221 mice to generate FVB Tg21221 MAPTnull mice homozygous for both thte Tg21221 transgene and the MAPT knockout. The other feeder line was generated by crossing B6C3 APP/PS1 mice with B6 MAPTnull mice to generate mice heterozygous for the APP/PS1 transgene and homozygous for the MAPT knockout. These two feeder lines were bred to generate F1 experimental animals. Human tau is only expressed when the tetracycline transactivator is also expressed and can be suppressed by feeding the mice doxycycline (figure 1A). This consistent outbred background breeding scheme keeps variability low while avoiding potential pitfalls of inbred strains such as sensory deficits during ageing, liver deficits, deletions such as loss of alpha-synuclein in some C57 strains, and other unknown recessive defects that may occur in inbred lines (Cudalbu et al., 2013; Specht and Schoepfer, 2001; Wong and Brown, 2006). Out of the 395 mice born during the generation and phenotyping of the MAPT-AD line, as expected 100% were homozygous for endogenous tau knockout, 100% were heterozygous for the rTg21221 tau responder transgene, 53% were heterozygous for the APP/PS1 transgene (50% expected), 48% were heterozygous for the CK-tTA activator transgene (50% expected), and 23% had both the APP/PS1 and CK-tTA transgenes (25% expected). 48% of the mice were female. Thus, the transgenes were all inherited in the expected Mendelian ratios, indicating that no combination of genotypes was lethal (Chi squared value = 6.41, p= 0.093, df = 3 confirming Mendalian ratios). This is an important advantage of our consistent outbred breeding scheme as the same APP/PS1 transgene is lethal to about half of the mice on a congenic B6 background (Bennett et al., 2017).

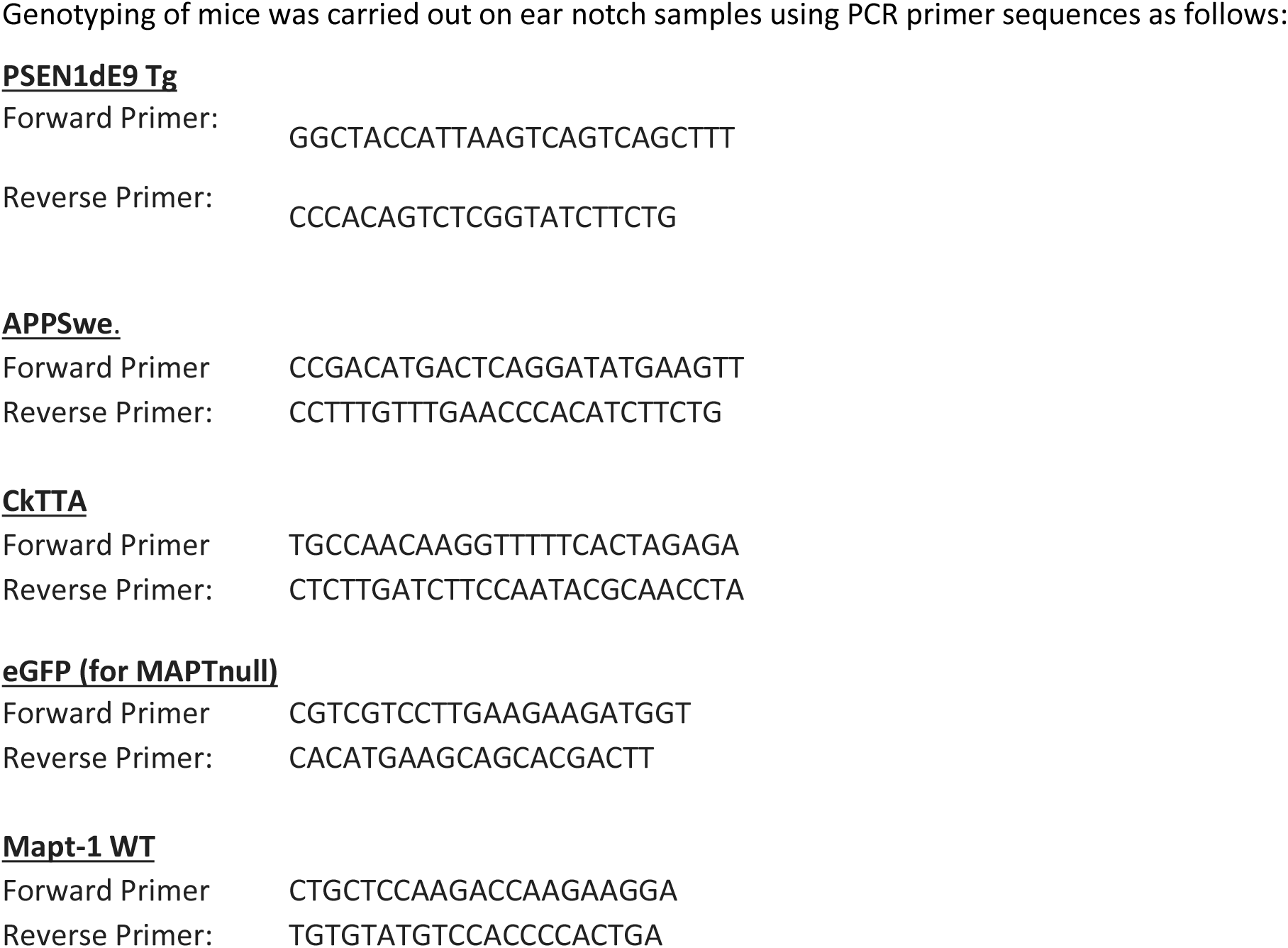

One cohort of mice was aged and used for behavioural testing at 3, 6, and 9-10 months of age and sacrificed at 9-10 months of age for LTP experiments and pathological characterization (see supplemental table 1 for all mouse data). Another cohort of mice was aged to 10-10.5 months of age, tested for baseline behaviour, then half of the mice were treated with 200ppm doxycycline in the chow for 4 months to reduce tau transgene expression and the others treated with vehicle. These mice were sacrificed at 14-14.5 months of age for pathological studies. Another cohort of littermates was aged to 6 months and sacrificed to look at onset of pathology.

As a negative control to be sure that any effects of tau expression were not an artefact of the CKtTA activator transgene, which is expressed in all mice that express tau by necessity, we examined B6.CKtTA mice on a mouse tau null background at 9 months of age for behavioural and pathological changes. As a positive control for the effects of the APP/PS1 transgene on behaviour and synaptic plasticity in our hands, we tested a 9-month-old cohort of B6C3.APP/PS1 mice from Jackson labs at 9 months of age. As a positive control for tau staining rTg4510 brain sections from 3 mice were used for tau immunohistochemistry (Santacruz et al., 2005; Spires et al., 2006).

#### Behavioural testing

Animals were tested for open field behaviour in a square box (40 x 40 x 60 cm) composed of dark opaque walls with approximately 2.5cm of corn cob bedding on the floor of the arena. Animals were recorded using an overhead camera and the video signal fed into Blackmagic Media Express computer software which captured the animals movements. Each day animals were brought into the testing room in their home cage upon the end of the 12 hr dark cycle and allowed to settle for 1 hour. For habituation, animals were exposed to the open field for 3 consecutive days. On day 1, animals were introduced to the centre of the arena along with cage mates for 20 minutes. For days 2-4, individual animals were placed facing a corner of the arena, which was assigned using a random generator. For each experimental group, the order in which animals were placed in the arena was randomly assigned using a random sequence generator. On day 4, behavior in the open field was recorded for 10 minutes using an overhead camera and movements captured with Blackmagic Media Express software. idTracker software and MATLAB were used to analyse mouse behaviour. The total distance travelled, distance travelled in the outer segment (40 x 40 – inner segment), distance travelled in the inner segment (20 x 20), percentage of time spent in the outer segment and percentage of time spent in the inner segment, were calculated and analysed in SPSS and Prism7.

In order to ensure that the hyperactivity observed at 14.5 months of age is not a consequence of baseline performance prior to treatment, 10.5 month old mice were assessed for baseline performance in the open field according to the treatment group to which they would be assigned. A significant effect of genotype was observed (p<0.0001), however there was no difference in open field behaviour in the cohorts destined for doxycycline or vehicle treatment within the same genotype (2-way ANOVA effect of treatment F(1,164)=0, p>0.99999). This suggests the increase in total distance travelled in 14.5 month old vehicle-treated MAPT-AD mice and reversal with doxycycline is not due to baseline increased activity in this group at 10.5 months of age.

#### Measuring pathology

Mice were sacrificed by terminal anaesthesia and perfused with PBS. Brains were dissected and one hemisphere fixed for 48 hours in 4% paraformaldehyde. Samples of entorhinal cortex from the other hemisphere were saved for array tomography as detailed below and the rest of the hemisphere was frozen for biochemical analyses. The fixed hemisphere was cryoprotected in 15% glycerol and sectioned into 50 micron coronal sections through the entire hemisphere with a sliding microtome (Leica SM2010R sliding microtome). To quantify amyloid pathology, every 20^th^ section was stained with a pan-Aβ antibody and counterstained with 0.05% Thioflavine S in 50% ethanol to label plaque fibrils and any neurofibrillary tangles (antibody details are found in table 2). Tile scan images of each entire section were obtained with a 10x objective on a Zeiss Axioimager microscope. Images were analysed using ImageJ. The cortex and hippocampus on each section were outlined, regions of interest defined, and the area calculated. Cortical and hippocampal volumes were estimated by multiplying the area on each section by 1000 (distance between sections), summing these values for all sections, and multiplying by 2 to estimate total volume as we only measured one hemisphere. Each channel of amyloid staining was manually thresholded in ImageJ by an experimenter blinded to genotype. The ImageJ analyze particles function was used to calculate the percent area of cortex and hippocampus occupied by staining and the number and average size of individual plaques. To calculate the burden of oligomeric halos surrounding plaques, the thresholded Thioflavin S image was subtracted from the thresholded pan-Aβ image and plaque burden, number, and size were analysed as above.

Series of every 10^th^ section were also stained with pathological tau antibodies to look for neurofibrillary tangles and neuropil threads as detailed in table 2. Stained sections were examined using a Zeiss AxioImager Z2 microscope and images acquired with a CoolSnap digital camera. For all immunostains, no primary conditions were used as negative controls. For tau stains, rTg4510 mouse brain sections containing neurofibrillary pathology were used as positive controls.

To measure gliosis, free floating coronal sections were stained for microglia (Iba1), astrocytes (GFAP), and fibrillary plaques (Thioflavine S), with citrate buffer pre-treatment (95°C for 20 minutes, see table 2 for antibody details). Three coronal sections were stained per mouse at approximately 0.75mm, - 2.0mm, and -3.75mm from Bregma. Tile scans were obtained at 10x magnification using a ZEISS Imager.Z2 stereology microscope and images were thresholded on ImageJ for cortical burden quantification.

#### RNA analyses

Total RNA was extracted from the frontal cortex using the Lipid Tissue Mini Kit (Qiagen). RNA quantity and quality was assessed using a Bioanalyzer 2100 (Agilent Technologies). All samples had RIN values > 7. To generate RNA-seq data, barcoded RNA-seq libraries were prepared by Edinburgh Genomics using the Illumina TruSeq stranded mRNA-seq kit, according to the manufacturer’s protocol (Illumina). The libraries were pooled and sequenced using an Illumina Novaseq 6000. RNA-sequencing was performed to a depth of ~60 million 50bp paired-end reads per sample. Reads were mapped to the mouse primary genome assembly (GRCm38) contained in Ensembl release 92 (Zerbino et al., 2018). Read alignment was performed with STAR (Dobin et al., 2013), version 2.5.3a, and tables of per-gene read counts were generated from the mapped reads with featureCounts (Liao et al., 2014), version 1.5.2. Differential expression analysis was then performed using DESeq2 (R package version 1.18.1) (Love et al., 2014). Gene Ontology enrichment analysis was performed with topGO [5] (R package version 2.30.1).

For qRT-PCR, cDNA was synthesised using the SuperScript VILO cDNA synthesis kit (ThermoFisher) and the following PCR settings used: 10 minutes at 25°C, 60 minutes at 42°C and 5 minutes at 85°C. qPCRs were run on a Stratagene Mx3000P QPCR System (Agilent Technologies) using SYBR Green MasterRox (Roche) with 6 ng of cDNA per well of a 96-well plate, using the following programme: 10 min at 95 °C, 40 cycles of 30 s at 95 °C, 40 s at 60 °C and 30 s at 72 °C, with a subsequent cycle of 1 min at 95 °C and 30 s at 55 °C ramping up to 95 °C over 30 s (to measure the dissociation curve). The following primers were used:

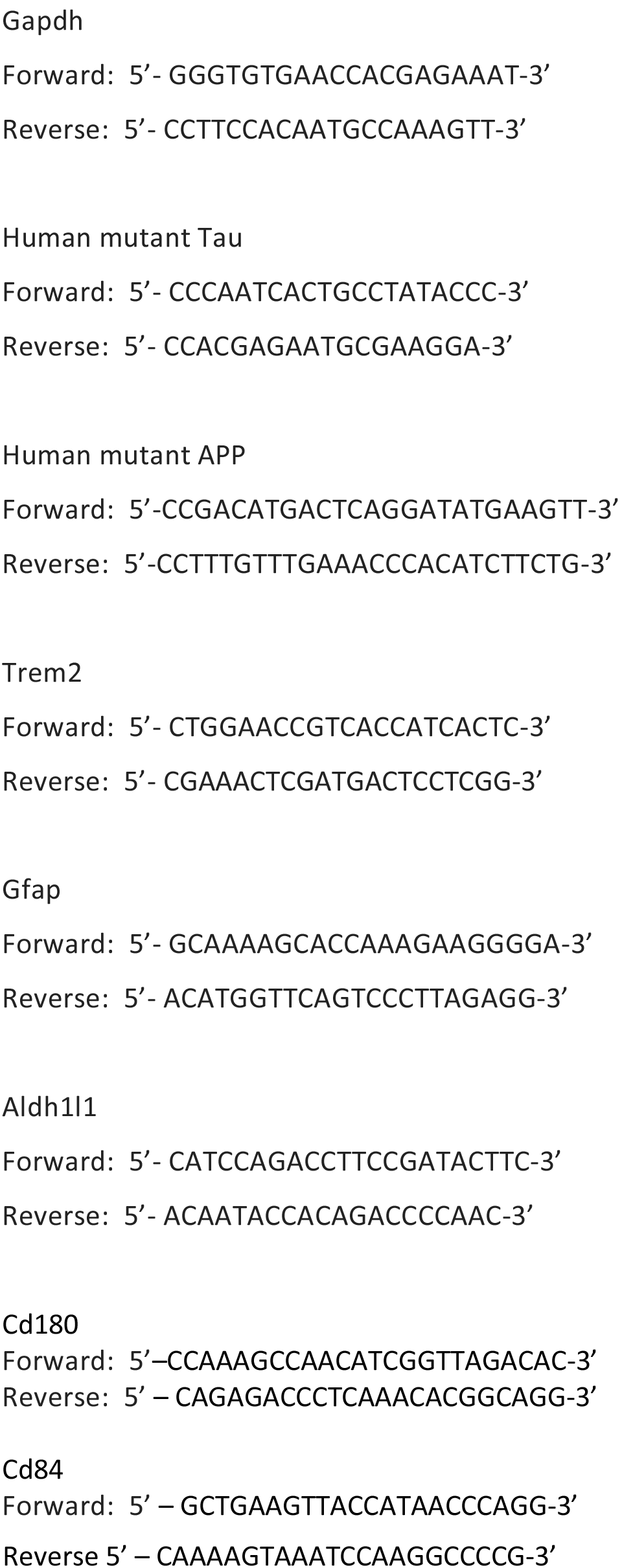

#### Array Tomography

Fresh brain tissue samples were collected from 14 month old mice and human subjects as outlined previously (Kay et al., 2013; Koffie et al., 2009). Small tissue blocks containing cortex were fixed in 4% paraformaldehyde and 2.5% sucrose in 20 mM phosphate buffered saline pH 7.4 (PBS) for 3 hours. Samples were then dehydrated through ascending cold graded ethanol and embedded into LR White resin (EMS) which was allowed to polymerise overnight at 53 °C. Resin embedded tissue blocks were cut into array ribbons of 70 nm thick sections using an ultracut microtome (Leica) equipped with a Jumbo Histo Diamond Knife (Diatome, Hatfield, PA) and collected onto gelatin coated coverslips.

For pathological protein colocalisation with post-synapses, array ribbons were immunostained with primary antibodies against total post synapses (PSD95), oligomeric amyloid-beta (1C22) and total tau (pan-tau). For pathological protein colocalisation with pre-synapses, array ribbons were immunostained with primary antibodies against total synaptic vesicle protein synaptophysin, amyloid-beta (AW7) and total tau (pan-tau) (Table 2). Sections were counterstained with 0.01 mg/mL 4’-6-diamidino-2- phenylindole (DAPI). In each experiment, a short extra ribbon was used as a no primary negative control. Images were obtained on serial sections using a Zeiss axio Imager Z2 epifluorescent microscope with a 10x objective for tile scans and 63x 1.4NA Plan Apochromat objective for high resolution images. Images were acquired with a CoolSnap digital camera and AxioImager software with array tomography macros (Carl Zeiss, Ltd, Cambridge UK).

Human brain Array tomography ribbons were stained with combinations of synaptic antibodies, tau antibodies and AW7 to label amyloid beta as described in the figures and table 2. For two-day stains, antibodies applied for the first imaging day were stripped by incubation in aqueous 0.02% SDS and 0.8% sodium hydroxide solution for 20 minutes. Stripped ribbons were rinsed in water and re-probed with another set of primary then secondary antibodies.

Images from each set of serial sections were converted into image stacks and aligned using the Image J plug-in, MultiStackReg (courtesy of Brad Busse and P. Thevenaz, Stanford University) (Thevenaz et al., 1998). Regions of interest within the cortical neuropil were chosen (10 μm^2^) and their proximity to plaque edges recorded (<20 μm from a plaque edge considered “near” plaques and >20 μm from a plaque edge considered “far” from plaques). Image stacks were then binarised using thresholding algorithms in ImageJ. For synaptic staining, images stacks were binarised using an ImageJ script that combines different thresholding algorithms in order to select both high and low intensity synapses in an automated and unbiased manner. To calculate the synaptic density, thresholded images were processed and analysed in MATLAB to remove background noise (objects present in only a single section were removed). To examine pathological protein presence at the synapse, thresholded images were processed and analysed in MATLAB to remove background noise and to calculate the colocalisation of total tau and oligomeric amyloid-beta with post synapses individually and in combination (a minimum of 50% of the synapse volume had to overlap with tau and/or 1C22 to qualify as positive for that stain).

### QUANTIFICATION AND STATISTICAL ANALYSIS

All experiments were carried out by a person blind to genotype and treatment of the mice and blind to diagnosis for human studies. For each experimental variable, a percentage, mean or median was calculated for each subject (mouse or human case). Groups of mice or people were compared with parametric or non-parametric tests as appropriate based on the normality of the datasets. Statistical tests were carried out in SPSS and Prism7. The number of subjects and statistical tests used for each experiment are indicated in the results, figure legends, and supplemental tables 1 and 2 (mouse) and 3 (human).

### DATA AND SOFTWARE AVAILABILITY

Spreadsheets of data used in this study are included as supplemental tables 1-3. All the RNA-seq data that support the findings of this study will be deposited in the European Bioinformatics Institute depository prior to publication. Custom imageJ and MATLAB macros used for image analysis are freely available on the University of Edinburgh Data Sharing repository.

**Figure S1 related to figure 1:**
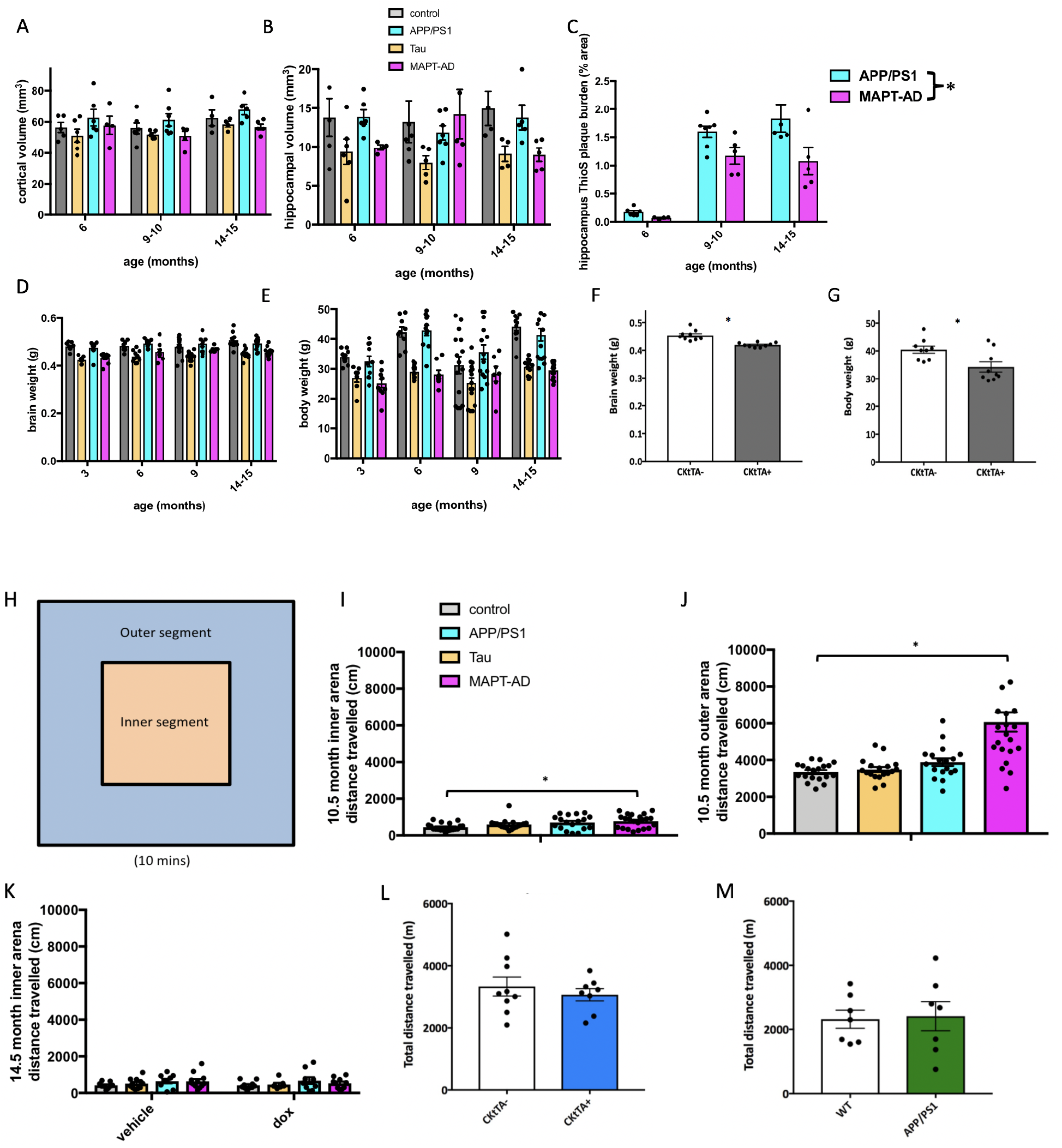
None of the genotypes experiences age related cortical (A) or hippocampal (B) atrophy. Hippocampal dense plaque burden is lower in MAPT-AD mice than APP/PS1 mice (C). Hippocampal volume is affected by genotype (2-way ANOVA F (3, 50) = 4.823, p=0.005) as are brain weight (D, F(3, 163) = 40.28, p<0.0001) and body weight (E, F(3, 163) = 37.25, p<0.0001). These effects are most likely driven by the CkTtA transgene that drives the tau responder (NOT tau expression) since the parent strain MAPTnullxCktTA mice have brain (F) and body weight (G) reductions at 9 months of age compared to MAPTnull littermate controls. To determine whether MAPT-AD mice have an anxiety phenotype, open field data was analysed by the distance travelled in the inner segment versus outer segment of the arena (H). At 10.5 months, there was a significant difference between genotypes in the inner (I, ANOVA, F[3,69]=4.075, p=0.010) and outer (J, ANOVA F[3,69]=15.91, p<0.0001) portions of the arena. MAPT-AD mice travelled significantly further in both the inner and outer arena compared to control mice (* Tukey’s posthoc test p<0.01). At 14.5 months, there were no significant differences between genotype and treatment in distance travelled in the inner arena (K). The hyperactivity phenotype in MAPT-AD mice is not driven by the CKtTA transgene that drives tau expression as mice with CKtTA only are no different from their littermate controls at 9-10 months of age (L). Similarly, the APP/PS1 mice with endogenous mouse tau do not have a hyperactivity phenotype at 9-10 months of age (M). Graphs depict mean ± SEM. Individual points represent the mean value for each mouse.

**Figure S2 related to figure 3:**
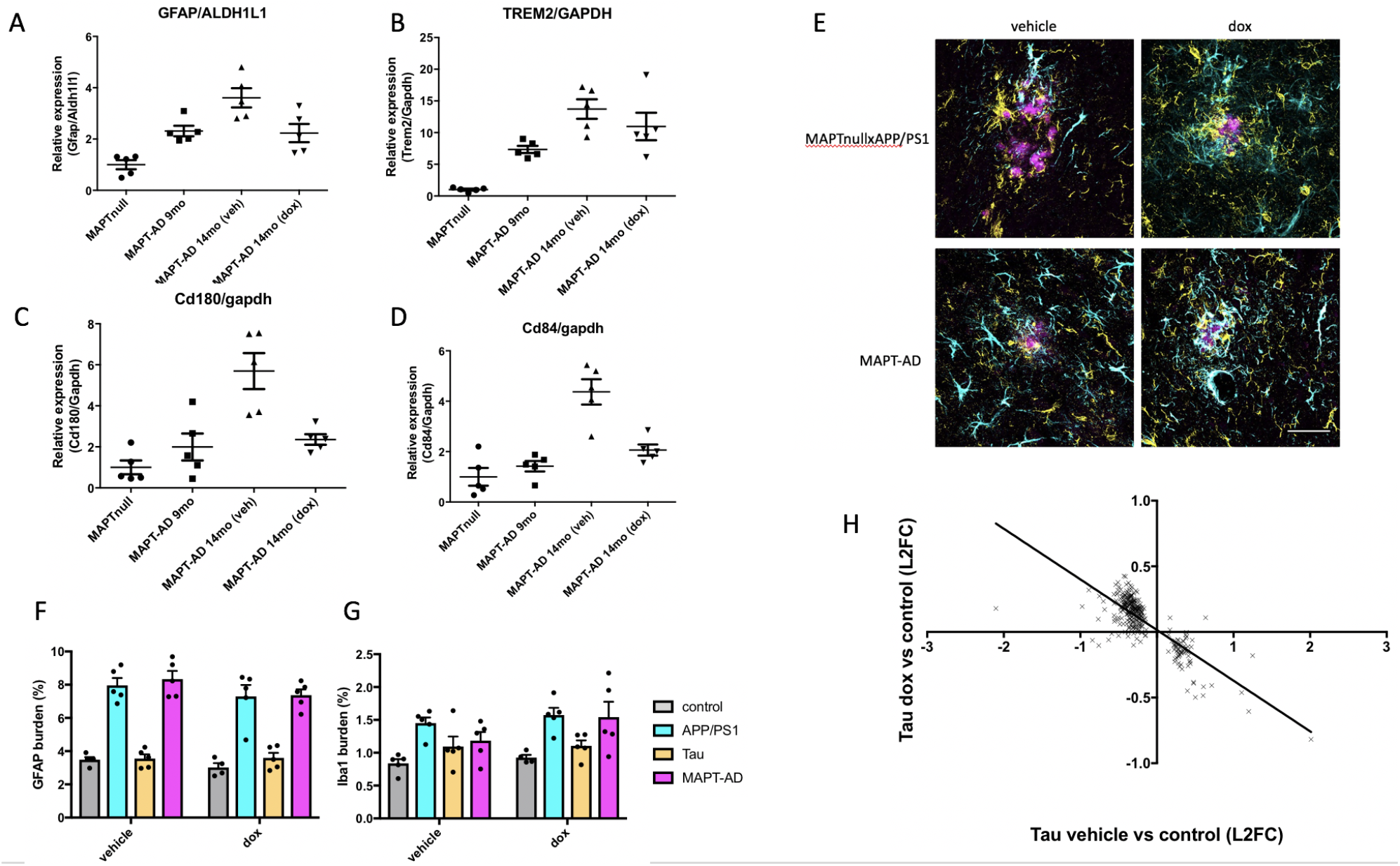
RT-PCR validation of RNAseq results at 9 months and 14.5 months of age indicate that the upregulated genes GFAP (A), Trem2 (B), Cd180 (C), and Cd84 (D) increase between 9 and 14.5 months and that this is prevented by dox treatment. The percentage area occupied by GFAP labelled astrocytes (Cyan, E) and Iba1 labelled microglia (magenta E) was higher in genotypes with plaques but did not change with tau transgene suppression (F, GFAP 2-way ANOVA effect of genotype F[3,31]=75.16, p<0.001, treatment F[1,31]=3.22, p=0.082, G, Iba1 2-qay ANOVA effect of genotype F[3.31]=9.05, p=0.0002, treatment F[1,31]2.48, p=0.13). Dox treatment significantly rescues transcriptional changes in Tau mice (H). Scale bar represents 40 μm.

## References

Barry, A.E., Klyubin, I., Mc Donald, J.M., Mably, A.J., Farrell, M.A., Scott, M., Walsh, D.M., and Rowan, M.J. (2011). Alzheimer’s disease brain-derived amyloid-beta-mediated inhibition of LTP in vivo is prevented by immunotargeting cellular prion protein. The Journal of neuroscience: the official journal of the Society for Neuroscience 31, 7259–7263.

Bennett, R.E., DeVos, S.L., Dujardin, S., Corjuc, B., Gor, R., Gonzalez, J., Roe, A.D., Frosch, M.P., Pitstick, R., Carlson, G.A., et al. (2017). Enhanced Tau Aggregation in the Presence of Amyloid beta. Am J Pathol 187, 1601–1612.

Crimins, J.L., Pooler, A., Polydoro, M., Luebke, J.I., and Spires-Jones, T.L. (2013). The intersection of amyloid beta and tau in glutamatergic synaptic dysfunction and collapse in Alzheimer’s disease. Ageing research reviews.

Cudalbu, C., McLin, V.A., Lei, H., Duarte, J.M., Rougemont, A.L., Oldani, G., Terraz, S., Toso, C., and Gruetter, R. (2013). The C57BL/6J mouse exhibits sporadic congenital portosystemic shunts. PloS one 8, e69782.

De Strooper, B., and Karran, E. (2016). The Cellular Phase of Alzheimer’s Disease. Cell 164, 603–615.

Dobin, A., Davis, C.A., Schlesinger, F., Drenkow, J., Zaleski, C., Jha, S., Batut, P., Chaisson, M., and Gingeras, T.R. (2013). STAR: ultrafast universal RNA-seq aligner. Bioinformatics 29, 15–21.

Fox, L.M., William, C.M., Adamowicz, D.H., Pitstick, R., Carlson, G.A., Spires-Jones, T.L., and Hyman, B.T. (2011). Soluble tau species, not neurofibrillary aggregates, disrupt neural system integration in a tau transgenic model. J Neuropathol Exp Neurol 70, 588–595.

Fu, H., Rodriguez, G.A., Herman, M., Emrani, S., Nahmani, E., Barrett, G., Figueroa, H.Y., Goldberg, E., Hussaini, S.A., and Duff, K.E. (2017). Tau Pathology Induces Excitatory Neuron Loss, Grid Cell Dysfunction, and Spatial Memory Deficits Reminiscent of Early Alzheimer’s Disease. Neuron 93, 533–541 e535.

Haas, L.T., and Strittmatter, S.M. (2016). Oligomers of Amyloid beta Prevent Physiological Activation of the Cellular Prion Protein-Metabotropic Glutamate Receptor 5 Complex by Glutamate in Alzheimer Disease. J Biol Chem 291, 17112–17112.

Hardy, J.A., and Higgins, G.A. (1992). Alzheimer’s disease: the amyloid cascade hypothesis. Science 256, 184–185.

Hong, S., Beja-Glasser, V.F., Nfonoyim, B.M., Frouin, A., Li, S., Ramakrishnan, S., Merry, K.M., Shi, Q., Rosenthal, A., Barres, B.A., et al. (2016). Complement and microglia mediate early synapse loss in Alzheimer mouse models. Science.

Hoover, B.R., Reed, M.N., Su, J., Penrod, R.D., Kotilinek, L.A., Grant, M.K., Pitstick, R., Carlson, G.A., Lanier, L.M., Yuan, L.-L., et al. (2010). Tau Mislocalization to Dendritic Spines Mediates Synaptic Dysfunction Independently of Neurodegeneration. Neuron 68, 1067–1081.

Hu, N.W., Corbett, G.T., Moore, S., Klyubin, I., O’Malley, T.T., Walsh, D.M., Livesey, F.J., and Rowan, M.J. (2018). Extracellular Forms of Abeta and Tau from iPSC Models of Alzheimer’s Disease Disrupt Synaptic Plasticity. Cell Rep 23, 1932–1938.

Hu, N.W., Nicoll, A.J., Zhang, D., Mably, A.J., O’Malley, T., Purro, S.A., Terry, C., Collinge, J., Walsh, D.M., and Rowan, M.J. (2014). mGlu5 receptors and cellular prion protein mediate amyloid-beta-facilitated synaptic long-term depression in vivo. Nat Commun 5, 3374.

Hudry, E., Wu, H.Y., Arbel-Ornath, M., Hashimoto, T., Matsouaka, R., Fan, Z., Spires-Jones, T.L., Betensky, R.A., Bacskai, B.J., and Hyman, B.T. (2012). Inhibition of the NFAT pathway alleviates amyloid beta neurotoxicity in a mouse model of Alzheimer’s disease. The Journal of neuroscience: the official journal of the Society for Neuroscience 32, 3176–3192.

Hyman, B.T. (2011). Amyloid-dependent and amyloid-independent stages of Alzheimer disease. Arch Neurol 68, 1062–1064.

Ittner, L.M., Ke, Y.D., Delerue, F., Bi, M., Gladbach, A., van Eersel, J., Wolfing, H., Chieng, B.C., Christie, M.J., Napier, I.A et al. (2010). Dendritic function of tau mediates amyloid-beta toxicity in Alzheimer’s disease mouse models. Cell 142, 387–397.

Izzo, N.J., Staniszewski, A., To, L., Fa, M., Teich, A.F., Saeed, F., Wostein, H., Walko, T., 3rd, Vaswani, A., Wardius, M et al. (2014). Alzheimer’s therapeutics targeting amyloid beta 1-42 oligomers I: Abeta 42 oligomer binding to specific neuronal receptors is displaced by drug candidates that improve cognitive deficits. PloS one 9, e111898.

Jackson, R.J., Rudinskiy, N., Herrmann, A.G., Croft, S., Kim, J.M., Petrova, V., Ramos-Rodriguez, J.J., Pitstick, R., Wegmann, S., Garcia-Alloza, M et al. (2016). Human tau increases amyloid beta plaque size but not amyloid beta-mediated synapse loss in a novel mouse model of Alzheimer’s disease. Eur J Neurosci 44, 3056–3056.

Jankowsky, J.L., Fadale, D.J., Anderson, J., Xu, G.M., Gonzales, V., Jenkins, N.A., Copeland, N.G., Lee, M.K., Younkin, L.H., Wagner, S.L., et al. (2004). Mutant presenilins specifically elevate the levels of the 42 residue beta-amyloid peptide in vivo: evidence for augmentation of a 42-specific gamma secretase. Hum Mol Genet 13, 159–170.

Jarosz-Griffiths, H.H., Noble, E., Rushworth, J.V., and Hooper, N.M. (2016). Amyloid-beta Receptors: The Good, the Bad, and the Prion Protein. J Biol Chem 291, 3174–3183.

Jay, T.R., Hirsch, A.M., Broihier, M.L., Miller, C.M., Neilson, L.E., Ransohoff, R.M., Lamb, B.T., and Landreth, G.E. (2017). Disease Progression-Dependent Effects of TREM2 Deficiency in a Mouse Model of Alzheimer’s Disease. The Journal of neuroscience: the official journal of the Society for Neuroscience 37, 637–637.

Kay, K.R., Smith, C., Wright, A.K., Serrano-Pozo, A., Pooler, A.M., Koffie, R., Bastin, M.E., Bak, T.H., Abrahams, S., Kopeikina, K.J., et al. (2013). Studying synapses in human brain with array tomography and electron microscopy. Nat Protoc 8, 1366–1366.

Keren-Shaul, H., Spinrad, A., Weiner, A., Matcovitch-Natan, O., Dvir-Szternfeld, R., Ulland, T.K., David, E., Baruch, K., Lara-Astaiso, D., Toth, B., et al. (2017). A Unique Microglia Type Associated with Restricting Development of Alzheimer’s Disease. Cell 169, 1276–1290 e1217.

Klein, W. (2013). Synaptotoxic amyloid-β oligomers: a molecular basis for the cause, diagnosis, and treatment of Alzheimer’s disease? Journal of Alzheimer’s disease: JAD 33 Suppl 1, 65.

Koffie, R.M., Hashimoto, T., Tai, H.C., Kay, K.R., Serrano-Pozo, A., Joyner1, D., Hou, S., Kopeikina, K.J., Frosch, M.P., Lee, V.M., et al. (2012). Apolipoprotein E4 effects in Alzheimer’s disease are mediated by synaptotoxic oligomeric amyloid-beta. Brain 135, 2155–2168.

Koffie, R.M., Meyer-Luehmann, M., Hashimoto, T., Adams, K.W., Mielke, M.L., Garcia-Alloza, M., Micheva, K.D., Smith, S.J., Kim, M.L., Lee, V.M., et al. (2009). Oligomeric amyloid beta associates with postsynaptic densities and correlates with excitatory synapse loss near senile plaques. Proc Natl Acad Sci U S A 106, 4012–4017.

Kopeikina, K.J., Hyman, B.T., and Spires-Jones, T.L. (2012). Soluble forms of tau are toxic in Alzheimer’s disease. Translational neuroscience 3, 223–223.

Kuchibhotla, K.V., Goldman, S.T., Lattarulo, C.R., Wu, H.Y., Hyman, B.T., and Bacskai, B.J. (2008). Abeta plaques lead to aberrant regulation of calcium homeostasis in vivo resulting in structural and functional disruption of neuronal networks. Neuron 59, 214–225.

Leyns, C.E.G., Ulrich, J.D., Finn, M.B., Stewart, F.R., Koscal, L.J., Remolina Serrano, J., Robinson, G.O., Anderson, E., Colonna, M., and Holtzman, D.M. (2017). TREM2 deficiency attenuates neuroinflammation and protects against neurodegeneration in a mouse model of tauopathy. Proc Natl Acad Sci U S A 114, 11524–11529.

Li, S., Hong, S., Shepardson, N.E., Walsh, D.M., Shankar, G.M., and Selkoe, D. (2009). Soluble oligomers of amyloid Beta protein facilitate hippocampal long-term depression by disrupting neuronal glutamate uptake. Neuron 62, 788–801.

Liao, Y., Smyth, G.K., and Shi, W. (2014). featureCounts: an efficient general purpose program for assigning sequence reads to genomic features. Bioinformatics 30, 923–930.

Love, M.I., Huber, W., and Anders, S. (2014). Moderated estimation of fold change and dispersion for RNA-seq data with DESeq2. Genome Biol 15, 550.

Lovestone, S., Hye, A., and Dixit, A. (2014). Clusterin as an early medator of Ab-induced disease processes: evidence from man Alzheimer’s & Dementia 10, 161.

Marzo, A., Galli, S., Lopes, D., McLeod, F., Podpolny, M., Segovia-Roldan, M., Ciani, L., Purro, S., Cacucci, F., Gibb, A., et al. (2016). Reversal of Synapse Degeneration by Restoring Wnt Signaling in the Adult Hippocampus. Curr Biol 26, 2551–2561.

Mattson, M.P., Cheng, B., Davis, D., Bryant, K., Lieberburg, I., and Rydel, R.E. (1992). beta-Amyloid peptides destabilize calcium homeostasis and render human cortical neurons vulnerable to excitotoxicity. The Journal of neuroscience: the official journal of the Society for Neuroscience 12, 376-389.

McInnes, J., Wierda, K., Snellinx, A., Bounti, L., Wang, Y.C., Stancu, I.C., Apostolo, N., Gevaert, K., Dewachter, I., Spires-Jones, T.L., et al. (2018). Synaptogyrin-3 Mediates Presynaptic Dysfunction Induced by Tau. Neuron 97, 823-835 e828.

Menkes-Caspi, N., Yamin, H.G., Kellner, V., Spires-Jones, T.L., Cohen, D., and Stern, E.A. (2015). Pathological tau disrupts ongoing network activity. Neuron 85, 959–966.

Mucke, L., and Selkoe, D. (2012). Neurotoxicity of amyloid β-protein: synaptic and network dysfunction. Cold Spring Harb Perspect Med 2.

Ovsepian, S.V., O’Leary, V.B., Zaborszky, L., Ntziachristos, V., and Dolly, J.O. (2018). Synaptic vesicle cycle and amyloid beta: Biting the hand that feeds. Alzheimers Dement 14, 502–513.

Prince, M., Anders, W., Guercget, M., Ali, G.-C., Wu, Y.-T., and Prina, M. (2015). World Alzheiemr Report 2015-the Global Impact of Dementia (Alzheimer’s Disease International).

Purro, S.A., Dickins, E.M., and Salinas, P.C. (2012). The secreted Wnt antagonist Dickkopf-1 is required for amyloid β-mediated synaptic loss. The Journal of Neuroscience.

Roberson, E.D., Halabisky, B., Yoo, J.W., Yao, J., Chin, J., Yan, F., Wu, T., Hamto, P., Devidze, N., Yu, G.Q., et al. (2011). Amyloid-beta/Fyn-induced synaptic, network, and cognitive impairments depend on tau levels in multiple mouse models of Alzheimer’s disease. The Journal of neuroscience: the official journal of the Society for Neuroscience 31, 700–711.

Roberson, E.D., Scearce-Levie, K., Palop, J.J., Yan, F., Cheng, I.H., Wu, T., Gerstein, H., Yu, G.Q., and Mucke, L. (2007). Reducing endogenous tau ameliorates amyloid beta-induced deficits in an Alzheimer’s disease mouse model. Science 316, 750–754.

Santacruz, K., Lewis, J., Spires, T., Paulson, J., Kotilinek, L., Ingelsson, M., Guimaraes, A., DeTure, M., Ramsden, M., McGowan, E et al. (2005). Tau suppression in a neurodegenerative mouse model improves memory function. Science 309, 476–481.

Sasaguri, H., Nilsson, P., Hashimoto, S., Nagata, K., Saito, T., De Strooper, B., Hardy, J., Vassar, R., Winblad, B., and Saido, T.C. (2017). APP mouse models for Alzheimer’s disease preclinical studies. EMBO J 36, 2473–2487.

Sellers, K.J., Elliott, C., Jackson, J., Ghosh, A., Ribe, E., Rojo, A.I., Jarosz-Griffiths, H.H., Watson, I.A., Xia, W., Semenov, M et al. (2018). Amyloid beta synaptotoxicity is Wnt-PCP dependent and blocked by fasudil. Alzheimers Dement 14, 306–317.

Shi, Q., Chowdhury, S., Ma, R., Le, K.X., Hong, S., Caldarone, B.J., Stevens, B., and Lemere, C.A. (2017a). Complement C3 deficiency protects against neurodegeneration in aged plaque-rich APP/PS1 mice. Sci Transl Med 9.

Shi, Y., Yamada, K., Liddelow, S.A., Smith, S.T., Zhao, L., Luo, W., Tsai, R.M., Spina, S., Grinberg, L.T.,Rojas, J.C., et al. (2017b). ApoE4 markedly exacerbates tau-mediated neurodegeneration in a mouse model of tauopathy. Nature 549, 523–527.

Shipton, O., Leitz, J., Dworzak, J., Acton, C., Tunbridge, E., Denk, F., Dawson, H., Vitek, M., Wade-Martins, R., Paulsen, O et al. (2011). Tau protein is required for amyloid {beta}-induced impairment of hippocampal long-term potentiation. The Journal of neuroscience: the official journal of the Society for Neuroscience 31, 1688–1692.

Small, S.A., and Duff, K. (2008). Linking Aβ and tau in late-onset Alzheimer’s disease: a dual pathway hypothesis. Neuron.

Specht, C.G., and Schoepfer, R. (2001). Deletion of the alpha-synuclein locus in a subpopulation of C57BL/6J inbred mice. BMC Neurosci 2, 11.

Spires, T.L., Meyer-Luehmann, M., Stern, E.A., McLean, P.J., Skoch, J., Nguyen, P.T., Bacskai, B.J., and Hyman, B.T. (2005). Dendritic spine abnormalities in amyloid precursor protein transgenic mice demonstrated by gene transfer and intravital multiphoton microscopy. The Journal of neuroscience: the official journal of the Society for Neuroscience 25, 7278–7287.

Spires, T.L., Orne, J.D., SantaCruz, K., Pitstick, R., Carlson, G.A., Ashe, K.H., and Hyman, B.T. (2006). Region-specific dissociation of neuronal loss and neurofibrillary pathology in a mouse model of tauopathy. Am J Pathol 168, 1598–1607.

Spires-Jones, T.L., Attems, J., and Thal, D.R. (2017). Interactions of pathological proteins in neurodegenerative diseases. Acta Neuropathol 134, 187–205.

Spires-Jones, T.L., and Hyman, B.T. (2014). The intersection of amyloid beta and tau at synapses in Alzheimer’s disease. Neuron 82, 756–771.

Spires-Jones, T.L., Meyer-Luehmann, M., Osetek, J.D., Jones, P.B., Stern, E.A., Bacskai, B.J., and Hyman, B.T. (2007). Impaired spine stability underlies plaque-related spine loss in an Alzheimer’s disease mouse model. Am J Pathol 171, 1304–1311.

Spires-Jones, T.L., Mielke, M.L., Rozkalne, A., Meyer-Luehmann, M., de Calignon, A., Bacskai, B.J.,Schenk, D., and Hyman, B.T. (2009). Passive immunotherapy rapidly increases structural plasticity in a mouse model of Alzheimer disease. Neurobiol Dis 33, 213–213.

Terry, R.D., Masliah, E., Salmon, D.P., Butters, N., DeTeresa, R., Hill, R., Hansen, L.A., and Katzman, R. (1991). Physical basis of cognitive alterations in Alzheimer’s disease: synapse loss is the major correlate of cognitive impairment. Ann Neurol 30, 572–580.

Thevenaz, P., Ruttimann, U.E., and Unser, M. (1998). A pyramid approach to subpixel registration based on intensity. IEEE Trans Image Process 7, 27–27.

Tucker, K.L., Meyer, M., and Barde, Y.A. (2001). Neurotrophins are required for nerve growth during development. Nat Neurosci 4, 29–29.

Um, J.W., Kaufman, A.C., Kostylev, M., Heiss, J.K., Stagi, M., Takahashi, H., Kerrisk, M.E., Vortmeyer, A., Wisniewski, T., Koleske, A.J., et al. (2013). Metabotropic glutamate receptor 5 is a coreceptor for Alzheimer abeta oligomer bound to cellular prion protein. Neuron 79, 887–887.

Um, J.W., Nygaard, H.B., Heiss, J.K., Kostylev, M.A., Stagi, M., Vortmeyer, A., Wisniewski, T., Gunther, E.C., and Strittmatter, S.M. (2012). Alzheimer amyloid-beta oligomer bound to postsynaptic prion protein activates Fyn to impair neurons. Nat Neurosci 15, 1227–1235.

Vargas-Caballero, M., Denk, F., Wobst, H.J., Arch, E., Pegasiou, C.M., Oliver, P.L., Shipton, O.A., Paulsen, O., and Wade-Martins, R. (2017). Wild-Type, but Not Mutant N296H, Human Tau Restores Abeta-Mediated Inhibition of LTP in Tau(-/-) mice. Front Neurosci 11, 201.

Wong, A.A., and Brown, R.E. (2006). Visual detection, pattern discrimination and visual acuity in 14 strains of mice. Genes Brain Behav 5, 389–389.

Wu, H.Y., Hudry, E., Hashimoto, T., Kuchibhotla, K., Rozkalne, A., Fan, Z., Spires-Jones, T., Xie, H., Arbel-Ornath, M., Grosskreutz, C.L., et al. (2010). Amyloid beta induces the morphological neurodegenerative triad of spine loss, dendritic simplification, and neuritic dystrophies through calcineurin activation. The Journal of neuroscience: the official journal of the Society for Neuroscience 30, 2636–2649.

Yasuda, M., and Mayford, M.R. (2006). CaMKII activation in the entorhinal cortex disrupts previously encoded spatial memory. Neuron 50, 309–318.

Yeh, F.L., Wang, Y., Tom, I., Gonzalez, L.C., and Sheng, M. (2016). TREM2 Binds to Apolipoproteins, Including APOE and CLU/APOJ, and Thereby Facilitates Uptake of Amyloid-Beta by Microglia. Neuron 91, 328–340.

Yin, Y., Gao, D., Wang, Y., Wang, Z.H., Wang, X., Ye, J., Wu, D., Fang, L., Pi, G., Yang, Y., et al. (2016). Tau accumulation induces synaptic impairment and memory deficit by calcineurin-mediated inactivation of nuclear CaMKIV/CREB signaling. Proc Natl Acad Sci U S A 113, E3773–3781.

Zempel, H., Thies, E., Mandelkow, E., and Mandelkow, E.-M. (2010). A{beta} Oligomers Cause Localized Ca2+ Elevation, Missorting of Endogenous Tau into Dendrites, Tau Phosphorylation, and Destruction of Microtubules and Spines. J Neurosci 30, 11938–11950.

Zerbino, D.R., Achuthan, P., Akanni, W., Amode, M.R., Barrell, D., Bhai, J., Billis, K., Cummins, C., Gall, A., Giron, C.G., et al. (2018). Ensembl 2018. Nucleic Acids Res 46, D754–D761.

Zhou, L., McInnes, J., Wierda, K., Holt, M., Herrmann, A.G., Jackson, R.J., Wang, Y.C., Swerts, J., Beyens, J., Miskiewicz, K et al. (2017). Tau association with synaptic vesicles causes presynaptic dysfunction.Nat Commun 8, 15295.

